# The DanRI regulatory system in uropathogenic *Escherichia coli* subverts neutrophil responses

**DOI:** 10.1101/2025.06.10.658900

**Authors:** Dora Čerina, Matthieu Rousseau, Carla Hart Olaiz, Gerben Marsman, Arturo Zychlinsky, Molly A. Ingersoll

## Abstract

Uropathogenic *Escherichia coli* (UPEC) is the primary cause of urinary tract infections (UTI). To establish an infection, UPEC must evade infiltrating neutrophils and their antimicrobial neutrophil extracellular traps (NETs). In this study, we identify a previously uncharacterized two-gene regulatory system within the pathogenicity island PAI_UTI89_II, which we named DanRI (<u>D</u>efense <u>a</u>gainst <u>n</u>eutrophil <u>R</u>egulator and <u>I</u>nhibitor). DanRI is induced by nucleosomes present in NETs and enables UPEC to suppress neutrophil responses by attenuating reactive oxygen species production and NET formation. Mechanistically, DanI functions as an antagonist to the transcriptional regulator DanR, thereby modulating key bacterial processes, including metabolic processes, flagellar biosynthesis, and stress response pathways. DanRI is required for UPEC fitness and long-term persistence in a mouse model of UTI. Taken together, our findings reveal DanRI as a novel regulatory system that promotes UPEC pathogenesis through immune evasion.

**Graphical abstract and model:** 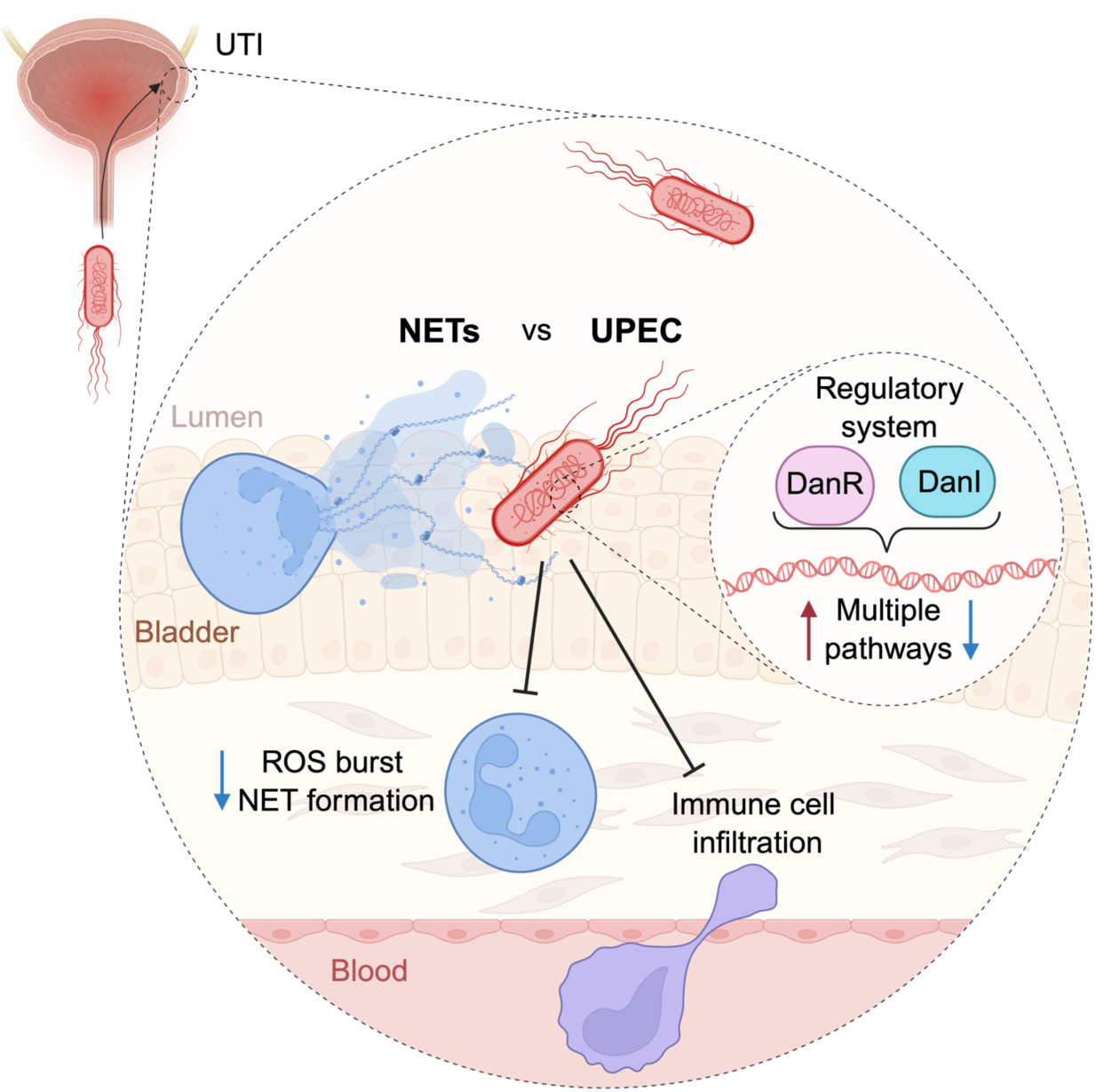

## Introduction

Uropathogenic *Escherichia coli* (UPEC) is responsible for approximately 80% of all urinary tract infections (UTI), a burden that disproportionately affects women of reproductive age.^1^ Recurrent infections are frequent, with almost half of affected individuals experiencing another episode within six months.^2,3^ While antibiotics remain the standard treatment, the growing prevalence of multidrug-resistant strains, particularly the globally disseminated *E. coli* sequence type (ST) 131, poses serious clinical challenges.^4,5^ With over 150 million UTI cases occurring annually worldwide and an increasing proportion attributed to multi-drug resistant UPEC, the impact of UTI is both substantial and growing.^6^ These trends highlight the urgent need to better define the molecular mechanisms that enable UPEC to persist and evade host immunity, to guide the development of targeted, antibiotic-independent therapies.

Pathogenic *E. coli* strains express virulence determinants that allow them to infect intestinal or extraintestinal niches, such as the urinary tract.^7^ During UTI, UPEC uses type 1 pili to ascend the urethra, adhere to and invade the urothelial lining, initiating infection within the bladder.^8^ The bladder responds to infection with cytokine and chemokine secretion, leading to robust neutrophil infiltration.^9^ As first responders of the innate immune system, neutrophils play a central role in bacterial clearance and coordination of the local immune response. In the bladder, they act not only to eliminate invading pathogens, but also engage other immune and stromal cells to restore homeostasis.^10–13^ To combat pathogens, neutrophils can phagocytose, degranulate, or form neutrophil extracellular traps (NETs).^14^

NETs are extracellular webs of chromatin adorned with cytosolic and nuclear proteins with antimicrobial properties. NETs entrap microbes extracellularly, concentrating effector proteins in the proximity of the microbe.^15^ Impaired formation of NETs leads to increased bacterial dissemination and exacerbation of UTI.^13^ NET components, including extracellular DNA, myeloperoxidase, and cathelicidin, are abundant in the urine of UTI patients.^16–18^ Interestingly, histones, the core structural elements of nucleosomes, are the most abundant proteins on NETs and potent antimicrobials.^19,20^ Other NET-associated effectors that can directly kill microbes include α-defensin, myeloperoxidase, and cathelicidin LL-37,^21–23^ while the protease neutrophil elastase can disarm microbes by cleaving their virulence factors.^15,24^

In the face of NET-mediated inhibition, pathogens have evolved strategies to inhibit NET formation or to actively degrade them.^25^ This counter-approach suggests that there is strong evolutionary pressure to evade and recognize this antimicrobial mechanism. Indeed, certain UPEC strains produce virulence factors, such as EsiB or YbcL,^26,27^ that can modulate neutrophil activation or migration, or TcpC, which might suppress NET production.^28^ These virulence factors are often encoded within pathogenicity islands or on the virulence plasmid,^7^ and their expression is coordinated by regulatory systems that respond to host-derived cues. While several such systems have been described in UPEC,^29^ it remains unknown whether, or by what mechanisms, UPEC detects and responds to NETs.

Here, we investigated the UPEC transcriptional response to NETs. We identified a previously uncharacterized two-gene regulatory system largely confined to UPEC strains, which we named DanRI (<u>D</u>efense <u>a</u>gainst <u>n</u>eutrophil <u>R</u>egulator and Inhibitor). The genes *danR* and *danI* are encoded within the pathogenicity island PAI_UTI89_II and are induced by NET-derived cues through distinct promoter elements. DanI inhibits the regulatory activity of DanR, modulating pathways such as flagellar synthesis, metabolic processes, and stress responses. Functionally, the DanRI system attenuated UPEC-induced neutrophil reactive oxygen species (ROS) burst and subsequent NET formation. Importantly, DanRI promoted UPEC fitness by dampening inflammation and facilitating immune evasion during bladder infection *in vivo*. Collectively, these findings establish DanRI as a key regulatory system shaping the interaction between UPEC and neutrophil responses during UTI.

## Results

### NETs stress UPEC and induce upregulation of uncharacterized genes

As NETs capture and eliminate microorganisms extracellularly,^15,30^ we assessed their antimicrobial activity on representative *E. coli* strains. We isolated neutrophils from the blood of healthy volunteers and stimulated them with phorbol 12-myristate 13-acetate (PMA). PMA is a widely used stimulus to activate protein kinase C and, like other mitogens, induces the formation of NETs.^31^ We incubated NETs with commensal *E. coli*, enteroinvasive *E. coli* (EIEC), enteropathogenic *E. coli* (EPEC), and UPEC, at a multiplicity of infection (MOI) of 10 for 1 hour, and quantified colony-forming units (CFU) to assess bacterial survival. NETS killed the commensal strain, but not the pathogenic strains of *E. coli* (**Figure 1A**), consistent with previous findings.^32^

**Figure 1.**
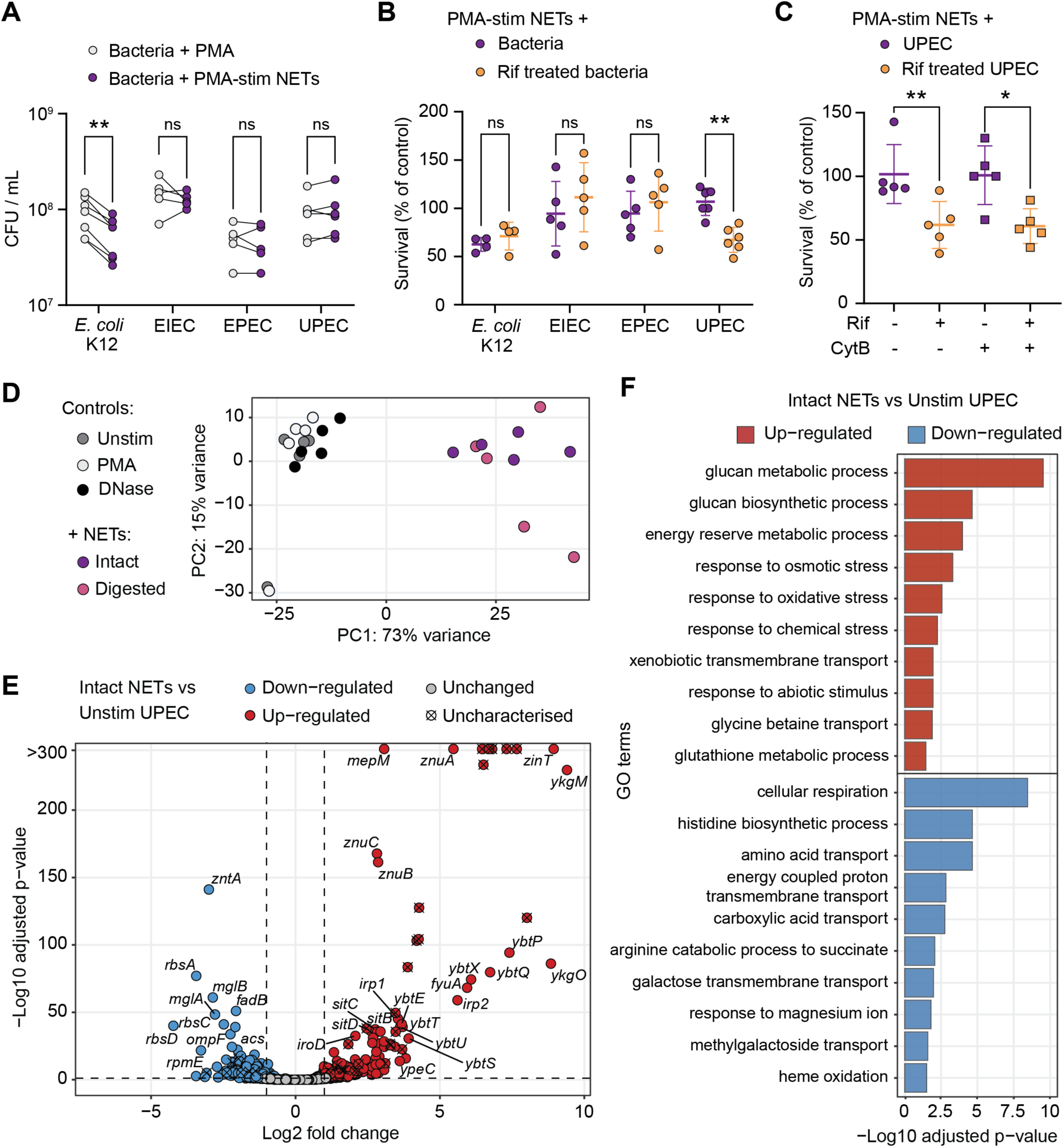
NETs upregulate stress responses and uncharacterized genes in UPEC. (A) Commensal *E. coli* K12, enteroinvasive *E. coli* (EIEC), enteropathogenic *E. coli* (EPEC), and uropathogenic *E. coli* (UPEC) were incubated with PMA-stimulated NETs at a MOI of 10 or in medium with PMA for 1 hour (n = 5-6). The graphs show data connected by lines to the matching control from the same experiment, and significance was determined with a paired t-test between control and stimulated condition (ns - no significant difference; ** p ≤ 0.01). (B) Commensal *E. coli* K12, EIEC, EPEC, and UPEC were treated with or without 10 µg/mL rifampicin to inhibit transcription before incubation with PMA-stimulated NETs at a MOI of 10 for 1 hour (mean ± SD, n = 4-6). Survival is expressed as a percentage of CFU after incubation with NETs, normalized to the CFU obtained from the medium with PMA or with rifampicin. Significance was determined by a Mann-Whitney test between control and stimulated condition (ns - no significant difference; ** p ≤ 0.01). (C) PMA-stimulated NETs were treated with or without 10 µg/mL cytochalasin B to inhibit phagocytosis. UPEC was treated with or without 10 µg/mL rifampicin to inhibit transcription and was added to NETs or medium with stimuli for 1 hour at a MOI of 10. Survival is expressed as a percentage of CFU of UPEC with NETs normalized to the appropriate controls of UPEC in medium with stimuli (mean ± SD, n = 5). Significance was determined by a Mann-Whitney test between control and stimulated condition (* p ≤ 0.05; ** p ≤ 0.01). (D) Principal component analysis of normalized gene expression of UPEC after 1 hour incubation in medium, with PMA, with DNase, and with PMA-stimulated NETs left intact or digested with DNase (n = 5). (E) Volcano plot showing differentially expressed genes in UPEC incubated with intact NETs compared to UPEC in medium alone. The graph depicts log2 fold change of gene expression plotted against the -log10 of the adjusted p-value. Downregulated genes are depicted in blue, unchanged genes in grey, upregulated genes in red, and uncharacterized differently expressed genes are depicted with an ‘x’. (F) Gene ontology enrichment analysis of upregulated genes depicted in red, and downregulated genes depicted in blue, in UPEC incubated with intact NETs compared to UPEC incubated in medium. Differentially expressed genes were associated with gene sets of *E. coli* K12 biological processes. The graph depicts the top 10 most significant terms of both categories and their affiliated -log10 of FDR-adjusted p-value.

Pathogenic bacteria encode virulence factors that enhance their survival in the presence of NETs.^25^ We hypothesized that upon interacting with NETs, *E. coli* upregulate evasion strategies to improve survival. To test the necessity of an active bacterial transcriptional response, we used the bacteriostatic RNA polymerase inhibitor rifampicin,^33^ which, as expected, reduced the total RNA levels across strains (**Figure S1A**). Interestingly, rifampicin-treated UPEC were more susceptible to killing by PMA-stimulated NETs compared to bacteria with intact transcription, a pattern not observed in other *E. coli* strains tested (**Figure 1B**). This suggested that UPEC initiates a transcriptional response to counter NETs. UPEC also required transcriptional activity to survive NETs when neutrophils were activated with nigericin, a molecule derived from *Streptomyces hygroscopicus*, that induces NETosis (**Figure S1B**).^32^ Additionally, to rule out the possibility that we were measuring phagocytosis-mediated killing, we added the phagocytosis inhibitor cytochalasin B and observed no change in antimicrobial activity, indicating that NETs alone were responsible for bacterial killing (**Figures 1C** and **S1B**).

We performed RNA-sequencing to investigate the transcriptional response of UPEC to NETs. We exposed UPEC to PMA-stimulated NETs that were either left intact or digested with DNase, and as controls, we assessed the effects of PMA and DNase alone on UPEC. The principal component analysis (PCA) revealed distinct gene expression changes in UPEC incubated with NETs (**Figure 1D**). Notably, the transcriptomic response activated by NETs was similar regardless of whether they were intact or digested, as the two conditions clustered closely together and the differentially expressed genes (DEGs) were correlated between the two conditions (**Figures 1D** and **S2A**). This suggests that UPEC responds to NET components and not necessarily the physical structures of NETs. Transcriptomic analysis revealed 390 upregulated and 269 downregulated genes in UPEC upon interaction with intact NETs compared to incubation in medium alone (**Figure 1E** and **Data S1**). Highly upregulated genes included those associated with responses to zinc starvation (*ykgM, ykgO, zinT, znuA, znuB, znuC*), siderophore systems including yersiniabactin (*ybtP, ybtQ, ybtX, ybtS, fyuA, irp1, irp2, ybtU, ybtT, ybtE*) and salmochelin (*iroB, iroC, iroD*), acid resistance (*hdeA, hdeB, gadE, gadX*), and oxidative stress response pathways (*msrP, msrQ*) (**Figure S2B**). The most strongly downregulated genes included those primarily involved in nutrient transport (*rbsA, rbsC, rbsD, mglA, mglB, dctA*) and energy production (*sdhA, sdhB, sdhC, sdhD, fdoI, fdoH, fadB*) (**Figure S2C**).

We performed gene ontology (GO) analysis^34^ to categorize all 659 DEGs in UPEC following NET exposure based on groups of biological processes. Upregulated genes were primarily linked to glucan metabolism and responses to osmotic, oxidative, and chemical stress, whereas downregulated genes were enriched for functions related to energy production and metabolite transporters (**Figure 1F**). The association of these terms and their underlying pathways suggests that UPEC adapts to the nutrient-limited environment imposed by NETs by mitigating stressors and preserving metabolic and energy resources. In addition to these pathways, a third of highly upregulated genes lacked functional characterization (**Figure S2B**), highlighting potentially unexplored aspects of UPEC responses to NETs.

### Uncharacterized genes upregulated by NETs are enriched in UPEC isolates

To investigate regulatory factors underlying the UPEC response to NETs, we analyzed the most highly upregulated genes for predicted DNA-binding motifs. This approach revealed a strongly induced, previously uncharacterized gene, UTI89_RS23725 (also known as UTI89_C4899; protein ID: WP_000841005.1), encoding a 122-amino-acid protein predominantly composed of a helix-turn-helix LytTR-type domain spanning approximately 100 amino acids. This domain is typically associated with transcriptional regulators of Gram-positive bacteria, where it functions in DNA binding and the regulation of virulence-associated gene expression.^35,36^ Immediately downstream of UTI90_RS23725, we identified two additional NET-induced loci: a hypothetical protein-coding gene, UTI89_RS27595 (protein ID: WP_001298030.1), and a pseudogene UTI89_RS23735, annotated as a transposase (**Figure 2A**). UTI89_RS27595 encodes a 69-amino-acid protein predicted to contain a domain of unknown function (DUF2116), characterized by a Zn-ribbon-like fold. All three genomic hits are situated within the pathogenicity island PAI_UTI89_II of the UPEC UTI89 genome (**Figure 2A**), integrated at the *leuX* chromosomal hotspot.^37,38^ This island also harbors several well-characterized virulence determinants, including P fimbriae, cytotoxic necrotizing factor 1, and hemolysin A.

**Figure 2.**
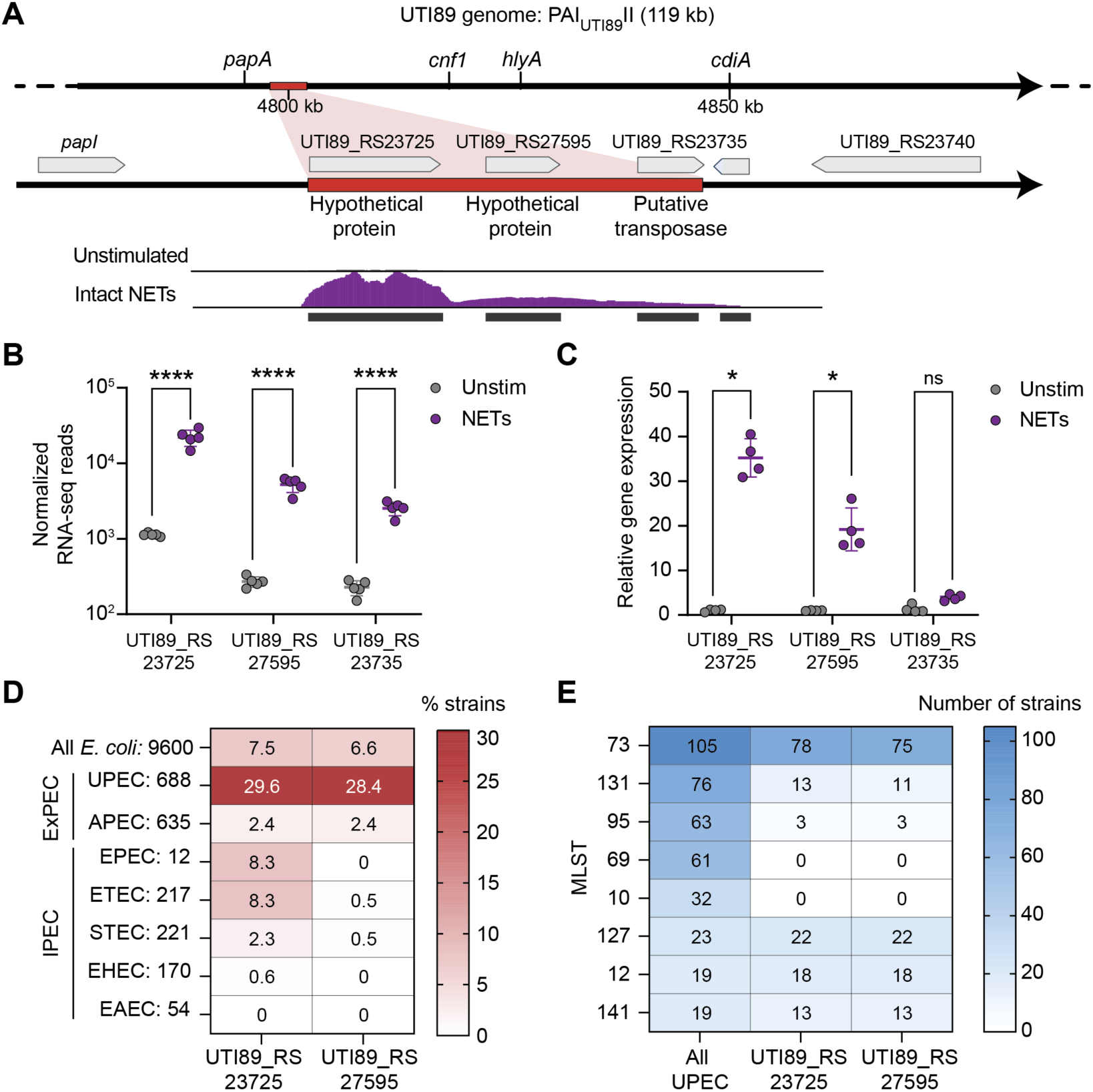
NETs upregulate uncharacterized genes enriched in UPEC genomes. (A) Graphical representation of the pathogenicity island PAI_UTI89_II with virulence factors (e.g., *papA, cnf1, hlyA,* and *cdiA*) and a zoom-in to the genomic region upregulated by NETs (in red). Representative transcript profiles of the genomic region in the UPEC in medium and UPEC with intact NETs. (B) RNA-sequencing read counts of the three genomic elements (UTI89_RS23725, UTI89_RS27595, and UTI89_RS23735) in UPEC exposed to intact NETs compared to UPEC in medium, normalized with the median of ratios method (mean ± SD, n = 5). Significance was determined with DESeq2 (**** p ≤ 0.0001). (C) Gene expression measured by RT-qPCR of three genomic elements (UTI89_RS23725, UTI89_RS27595 and UTI89_RS23735) in UPEC with NETs normalized to UPEC in medium for 1 hour relative to the housekeeping gene *gyrB* (mean ± SD, n = 4). Significance was determined by a Mann-Whitney test between control and stimulated condition (ns - no significant difference; * p ≤ 0.05). (D and E) Sequences of the two genes (UTI89_RS23725 and UTI89_RS27595) were aligned using the TBLASTX tool in the PubMLST database against the genomes of *E. coli*, which included extraintestinal pathogenic *E. coli* (ExPEC) strains: UPEC and avian pathogenic *E. coli (*APEC); and intestinal pathogenic *E. coli* (IPEC): Shiga toxin-producing (STEC), enterotoxigenic (ETEC), enterohemorrhagic (EHEC), enteroaggregative (EAEC), and enteropathogenic (EPEC) *E. coli.* The number of analyzed genomes is depicted next to the pathotype name. Strains with more than 50% identity were considered positive. The results of the sequence distribution over all pathotypes are plotted as a percentage of positive strains from the analyzed group (D). 688 UPEC strains were partitioned based on multilocus sequence typing (MLST). The graph depicts the number of all UPEC strains and sequence-positive strains for the two genes (UTI89_RS23725 and UTI89_RS27595) in the 8 largest MLST groups (E).

Transcriptome analysis revealed strong induction of the two hypothetical protein-coding genes UTI89_RS23725 and UTI89_RS27595, following NET exposure (**Figure 2B**), which was further validated by RT-qPCR (**Figure 2C**). By contrast, the adjacent pseudogene did not show significant upregulation (**Figure 2C**) and was therefore excluded from further analysis. Based on these findings, we focused subsequent investigations on the two NET-responsive hypothetical genes.

To identify homologous proteins, we performed NCBI protein BLAST searches^39^ and selected hits with more than 50% sequence identity. Closely related homologues of both proteins were found in *E. coli* strains. More distantly related homologues of UTI89_RS23725 were identified in *Salmonella enterica*, and in two *Yersinia* species (**Figures S3A** and **S3B**); while homologues of UTI89_RS27595 were also present in *S. enterica* strains (**Figures S3C** and **S3D**). All identified homologues encoded proteins of uncharacterized function.

Interestingly, using FoldSeek,^40^ we found that the protein encoded by UTI89_RS27595 shared structural motifs with two *E. coli* proteins: the RNA polymerase-binding transcription factor DksA,^41^ and the DNA gyrase inhibitor YacG.^42^ Both DksA and YacG interact with DNA-binding proteins to modulate their function. The region of similarity across these proteins includes a conserved four-cysteine Zn^2+^-binding region that forms a characteristic zinc finger domain, which is critical for their regulatory activity (**Figure S3D**).^43,44^

Given that these proteins are predominantly found in *E. coli*, we investigated their distribution across different *E. coli* pathotypes and clonal lineages. We analyzed 9,600 *E. coli* genomes from the PubMLST database.^45^ Using the TBLASTX against the translated nucleotide sequences, we selected hits with more than 50% sequence identity. We identified both genes in approximately 7% of all *E. coli* strains (**Figure 2D**). Among all *E. coli*, 1,997 strains were annotated with an associated pathotype, enabling assessment of gene prevalence across pathotypes. Strikingly, the genes were present in an average of 29% of UPEC strains but were rarely detected or absent in other *E. coli* pathotypes. When present, the two genes consistently displayed the same genomic arrangement, which is maintained according to their orientation, whether forward or reverse (**Figure S3E**), suggesting that, as expected in a pathogenicity island, they were co-inherited into the UPEC genome.

To investigate the gene distribution within the UPEC pathotype, we examined clonality using Achtman multilocus sequence typing. Most UPEC genomes harboring genes of interest belonged to ST73, the most prevalent clonal complex in our dataset and a clone frequently encountered in clinical settings (**Figure 2E**).^46^ Notably, the genes were also encoded in ST131, a widely disseminated multidrug-resistant UPEC clone.^4,5^ Additionally, these genes were prevalent in genomes associated with ST127, ST12, and ST141, less common UPEC clonal lineages that are increasingly associated with antibiotic resistance.^47–49^ The strong association of these two genes with clinical UPEC isolates and antibiotic resistance suggests a potential role in promoting UPEC pathogenesis.

### Autonomous promoters of *danR* and *danI* are activated by NET nucleosomes

Based on their predicted roles as components of a response regulatory system, we designated UTI89_RS23725 and UTI89_RS27595 as *danR* and *danI,* respectively – denoting “<u>d</u>efense <u>a</u>gainst <u>n</u>eutrophil <u>R</u>egulator” and “<u>d</u>efense <u>a</u>gainst <u>n</u>eutrophil <u>I</u>nhibitor”.

The genomic arrangement of *danR* and *danI* in *E. coli* genomes suggested that they may be organized as an operon. Operons typically consist of co-transcribed genes under a shared promoter,^50^ yet bacterial gene regulation is often more complex, especially under stress conditions, which can induce readthrough of adjacent genes.^51^ Our transcriptomic data revealed overlapping transcripts spanning *danR* and *danI* (**Figure 2A**), indicating they can be co-transcribed. However, the 139 bp intergenic region separating them is larger than expected for canonical operons and more characteristic of independently regulated genes.^50^ To resolve this ambiguity, we experimentally assessed the transcriptional architecture of the *danRI* locus.

We generated individual deletion mutants of *danR* and *danI*, as well as of the entire *danRI* region (**Figure 3A**), using the λ-Red recombination system.^52,53^ We complemented the mutants with either an empty pTrc99a plasmid (-p) or a plasmid carrying the deleted sequence (-*danRI*, -*danR*, -*danI*). All strains grew comparably in both rich LB medium and nutrient-limited human urine, indicating that the deletions did not impair general fitness (**Figure S4A**). RT-qPCR confirmed the absence of gene expression in deletion strains. Importantly, deletion of *danR* did not impact expression of *danI,* and vice versa, indicating that the two genes are independently regulated (**Figures 3B** and **3C**). Thus, the transcripts connecting *danR* and *danI* observed in our RNA-sequencing (**Figure 2A**) are likely a result of transcriptional readthrough triggered by NET-associated stress, as stressors can promote such events in bacteria.^51^ Finally, expression of the genes immediately upstream and downstream of *danRI* was unchanged in the Δ*danRI* strain compared to the wildtype (**Figure S4B**), indicating that the deletion did not exert a polar effect. We designed a luciferase-based reporter system to measure promoter activation. We selected the promoter sequences for *danR* and *danI* to encompass the majority of the regulatory sequence of each gene (**Figure 3A**). We cloned these sequences upstream from the *luxCDABE* operon, and the respective plasmids were introduced into UPEC. As expected, UPEC carrying a plasmid without a promoter or a plasmid with the original *em7* promoter showed no significant change in luciferase activity following 1 hour of NET exposure (**Figure 3D**). Notably, the promoters of *danR* and *danI* were significantly induced by NETs, indicating that these genes are independently activated via distinct, autonomous promoters (**Figure 3D**).

**Figure 3.**
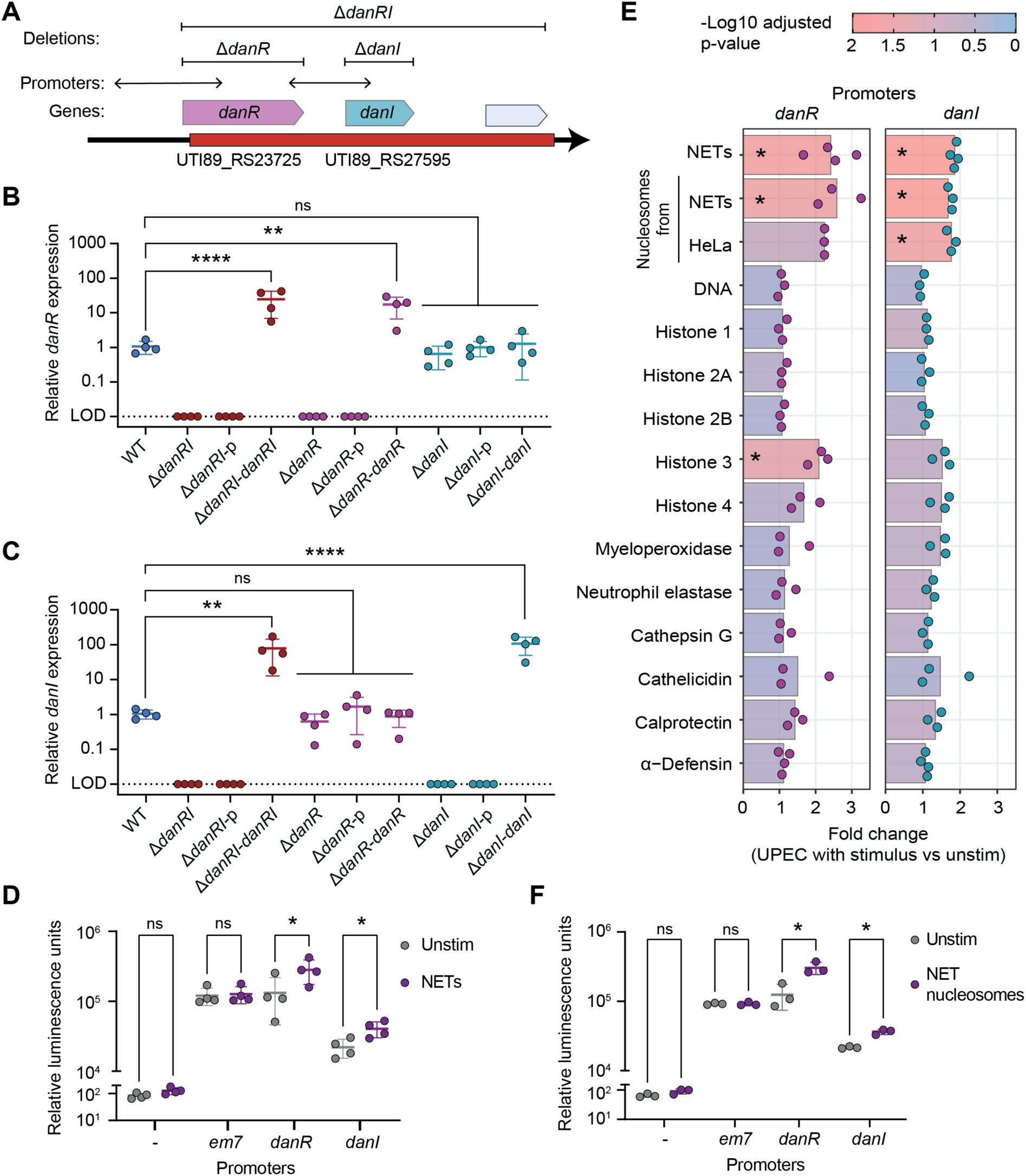
Nucleosomes from NETs activate *danR* and *danI* promoters. (A) Graphical representation of the genomic region of interest illustrates the *danR* and *danI* genes, along with the locations of targeted deletions and promoter sequences. (B and C) Gene expression measured by RT-qPCR of *danR* (B) and *danI* (C) in wildtype UPEC, deletion mutants, and their complemented strains relative to the housekeeping gene *gyrB* in medium (mean ± SD, n = 4). The limit of detection (LOD) is depicted as a dotted line. Significance was determined with one-way ANOVA with Šídák’s multiple comparisons test (ns - no significant difference; ** p ≤ 0.01; **** p ≤ 0.0001). (D, E, and F) UPEC reporter strains carrying plasmids with a luciferase operon under the control of no promoter (-), or promoter sequences of *em7*, *danR,* or *danI*, were incubated for 1 hour in medium with or without stimuli. Relative luminescence units were measured after incubation with PMA-stimulated NETs (D, mean ± SD, n = 4) or with nucleosomes purified from these NETs (F, mean ± SD, n = 3). Fold changes of luminescence units from unstimulated compared to stimulation with 15 different components in UPEC *danR* and *danI* reporter strains (E, n = 3-4). Significance was determined on luminescence values with a paired t-test between control and stimulated condition, corrected for the FDR (ns - no significant difference; * p ≤ 0.05). The –log10 of the FDR-adjusted p-value is represented by the color of the bar (E).

We tested the activation of the luciferase reporter strains by 15 different components of NETs, including DNA, nucleosomes, histones, and granular proteins (**Figure 3E**). We calculated the fold change in the luciferase signal and found that nucleosomes, both NET- and HeLa cell-derived, were strong activators of transcription, followed by histone H3. Notably, naked DNA, histones H1, H2A, H2B, H4, and several granular proteins did not stimulate the *danR* and *danI* promoters. NET-purified nucleosomes strongly activated the expression of both promoters, but not the *em7* promoter or the plasmid without a promoter (**Figure 3F**). These findings indicate that nucleosomes are the primary NET-derived signals activating the upregulation of the DanRI regulatory system.

## DanI inhibits DanR regulatory function across multiple pathways

Based on structural and sequence similarities, we hypothesized that DanR functions as a transcriptional regulator through the LytTR DNA-binding domain, while DanI regulates DanR activity. To assess their roles in gene regulation, we performed RNA-sequencing of wildtype, Δ*danRI*, Δ*danR,* and Δ*danI* strains under uninduced conditions. PCA revealed significant transcriptional changes in Δ*danI* compared to the wildtype UPEC strain, but minimal changes in the transcriptome of Δ*danRI* and Δ*danR* (**Figure 4A**). In the transcriptome of Δ*danRI* and *ΔdanR,* aside from the deleted genes, the only observed difference compared to the wildtype strain was the downregulation of the *ymgA* gene in *ΔdanR* (**Data S2**). Interestingly, the Δ*danI* transcriptome exhibited 122 upregulated and 287 downregulated DEGs (**Figure 4B** and **Data S2**). We used GO analysis on the 409 DEGs regulated in Δ*danI* and found that the upregulated genes were involved in many metabolic processes, while downregulated genes were primarily associated with flagellar assembly and organization (**Figure 4C**). The most strongly upregulated genes were associated with nutrient transport (*rbsA, rbsB, rbsC*), metabolic pathways (*rpiA*, *purC, purD*, *ldhA*, *gadB*, *guaC*, *fdaB*), and stress responses, such as acid resistance (*hdeA, hdeB, hdeD*), envelope stress (*pspA, pspB, pspC, pspG*), heat shock (*ibpB*), and starvation (*dsp*) (**Figure S5A**). The downregulated genes included those involved in flagellar assembly (*fliL, fliQ, fliF, fliP, fliH, fliO, fliM, fliJ, fliN, fliR*), S-fimbriae (*sfaA, sfaD, sfaE*), and catecholate siderophores receptors (*fiu, cirA, fepA*) (**Figure S5B**).

**Figure 4.**
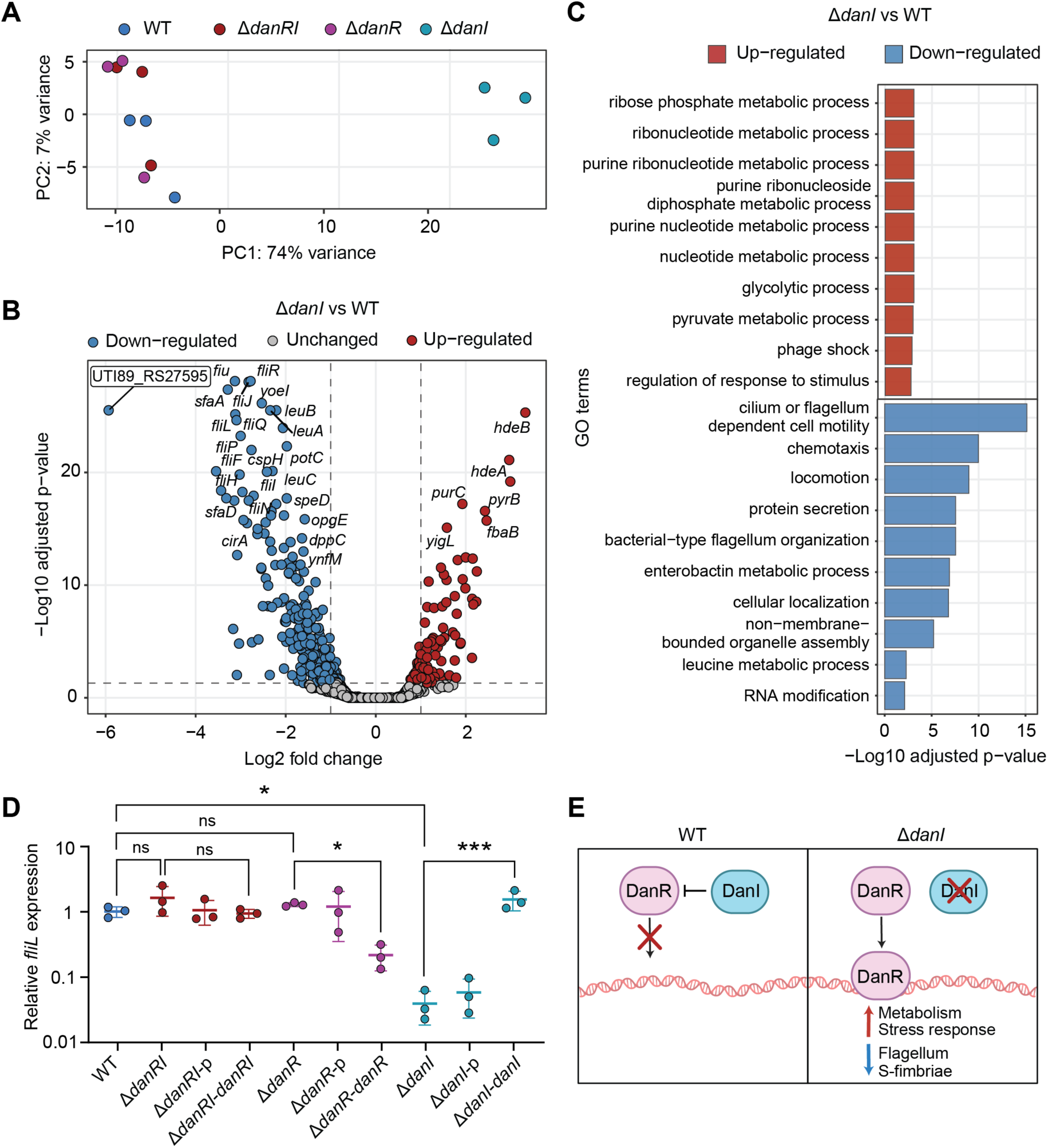
Deletion of *danI* changes gene expression. (A) Principal component analysis of normalized gene expression of the wildtype UPEC, Δ*danRI*, Δ*danR,* and Δ*danI* in media (n = 3). (B) Volcano plot showing differentially expressed genes in Δ*danI* compared to wildtype UPEC. The graph displays a log2 fold change of gene expression against the -log10 of the adjusted p-value. Downregulated genes are shown in blue, unchanged genes in grey, and upregulated genes in red. *DanI* (UTI89_RS27595) is highlighted. (C) Gene ontology enrichment analysis of differently expressed genes in Δ*danI* compared to the wildtype UPEC strain, using the biological processes in the *E. coli* K12 database. The graphs display the top 10 most significant terms of the upregulated genes, shown in red, and downregulated genes, in blue, along with their affiliated -log10 of FDR-adjusted p-values. (D) Gene expression measured by RT-qPCR of *fliL* gene in wildtype UPEC, deletion mutants, and their complemented strains relative to the housekeeping gene *gyrB* (mean ± SD, n = 3). Significance was determined with one-way ANOVA with Šídák’s multiple comparisons test (ns no significant difference; * p ≤ 0.05; *** p ≤ 0.001). (E) Graphical representation of the DanRI system regulation in wildtype UPEC and Δ*danI*

Intriguingly, while deletion of *danI* led to substantial transcriptional changes, the simultaneous deletion of both *danR* and *danI* in Δ*danRI* did not affect gene expression, suggesting a functional interplay between the two genes. Consistent with transcriptomic data, we found that *fliL* expression remained unchanged in Δ*danRI* and Δ*danR* but decreased in Δ*danI,* in comparison to the wildtype UPEC (**Figure 4D**). Complementation of *danI* in the Δ*danI* background restored *fliL* expression to wildtype UPEC levels. Interestingly, overexpression of *danR* in Δ*danR* reduced *fliL* expression, whereas co-expression of both genes in Δ*danRI* had no effect. These findings suggest that deletion of *danI* may relieve repression of *danR*, triggering the observed transcriptional changes. Overexpression of *danR* alone had a similar effect, reinforcing the idea that DanI functions as an inhibitor of DanR. Together, these data support a model in which DanI and DanR operate as a regulatory system, with DanR driving transcription and DanI modulating DanR activity (**Figure 4E**).

### The DanRI system dampens ROS and NET formation *in vitro*

NETs induced the upregulation of the DanRI regulatory system, which controlled the expression of several pathways, such as metabolic processes, flagellar biosynthesis, and stress responses. To assess whether DanRI influenced UPEC-neutrophil interactions, we examined the effects of its deletion on phagocytosis, ROS production, NETs release, and NET-mediated killing. Interestingly, Δ*danRI*, Δ*danR*, and Δ*danI* exhibited survival comparable to the wildtype UPEC strain when exposed to NETs (**Figure S6A**). As a positive control, we showed that Δ*neu*, a UPEC strain defective in capsule formation,^54^ was sensitive to NET-mediated killing (**Figure S6B**), confirming the importance of the capsule to protect against NETs.^55^ We incubated UPEC strains with live neutrophils for 30 minutes, then lysed the cells and enumerated bacterial CFU. As a control, bacteria were incubated in media alone or with neutrophils in the presence of cytochalasin B, with survival expressed as a percentage of CFU relative to the controls. Neutrophils killed wildtype UPEC, Δ*danRI*, Δ*danR,* and Δ*danI* comparably (**Figure S6C**), whereas Δ*neu* was also efficiently eliminated through phagocytosis (**Figures S6D** and **S6E**). These findings indicate that the DanRI regulatory system does not influence UPEC survival during NET-mediated killing or phagocytosis by neutrophils.

Upon neutrophil activation, the enzyme complex nicotinamide adenine dinucleotide phosphate (NADPH) oxidase generates a ROS burst.^14^ We measured real-time ROS production in neutrophils in response to infection with UPEC mutants or PMA for 3 hours. As expected, PMA induced a strong oxidative burst (**Figure 5A**).^31^ As a control, we showed that ROS production was inhibited by the addition of the NADPH oxidase inhibitor DPI (**Figure 5B**).^56^ Interestingly, the Δ*danRI* mutant induced a stronger ROS burst than wildtype UPEC, and this effect decreased when we complemented the deletion (**Figures 5A** and **5B**). We also observed elevated ROS levels in Δ*danR* and Δ*danI* infection, which were reduced upon complementation (**Figure 5C**). These results suggest that the DanRI system modulates and suppresses the ROS burst intensity in neutrophils upon interaction with UPEC.

**Figure 5.**
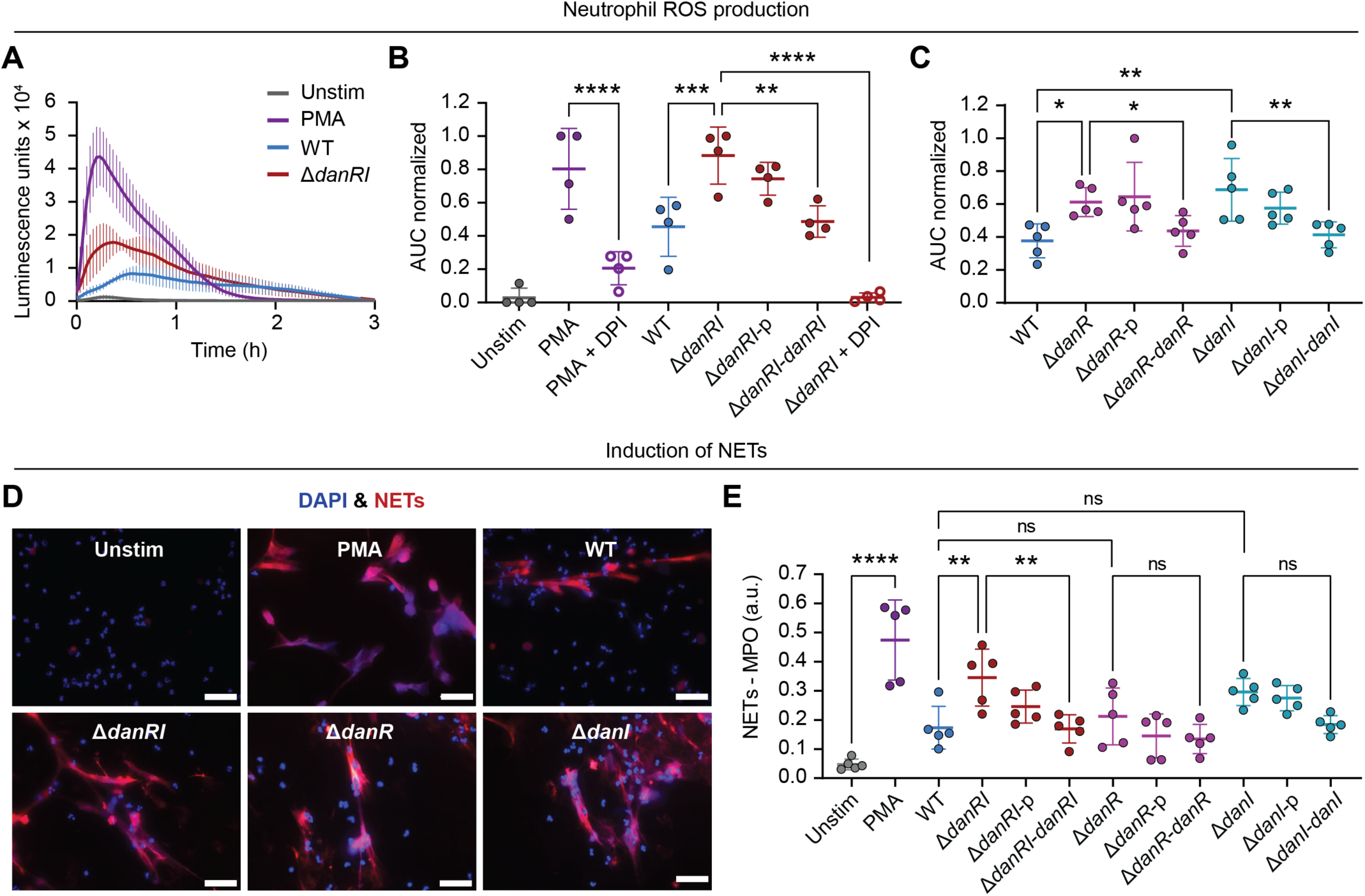
The regulatory system DanRI modulates neutrophil functions. (A, B, and C) Total reactive oxygen species (ROS) production was measured in neutrophils incubated with wildtype UPEC, deletion mutants, and their complemented strains, PMA, or just medium, for 3 hours using luminol chemiluminescence. Neutrophils were treated with 1 µM diphenyleneiodonium (DPI) to inhibit ROS production. Luminescence units are shown as ROS curves (A, mean ± SEM, n = 4). The area under the ROS curve (AUC) was calculated and normalized (B and C, mean ± SD, n = 4-5). (D and E) Neutrophils were incubated with wildtype UPEC, deletion mutants, complemented strains, PMA or medium for 4 hours, and NETs were visualized by immunofluorescence (D) or quantified (E). Representative images are shown for unstimulated, PMA, wildtype UPEC, Δ*danRI*, Δ*danR,* and Δ*danI*-stimulated NETs stained with DAPI (DNA, in blue) and 3D9 antibody detected with Alexa Fluor 647 (NETs, in red). The scale bar represents 50 µm (D). NETs were quantified with an ELISA using anti-MPO and 3D9 antibody (E, mean ± SD, n = 5). For all data, significance was determined using one-way ANOVA with Šídák’s multiple comparisons test (ns - no significant difference; * p ≤ 0.05; ** p ≤ 0.01; *** p ≤ 0.001; **** p ≤ 0.0001).

NET formation is reliant on the ROS burst.^32^ To determine whether the UPEC mutants affected NET induction, we stimulated neutrophils for 4 hours with the bacterial strains or with PMA, as a positive control. NETs were assessed microscopically (**Figure 5D**) and quantified by ELISA (**Figure 5E**), using a NET-specific antibody.^57^ PMA induced robust NET formation, and consistent with ROS measurements, the Δ*danRI* mutant triggered significantly more NETs than wildtype UPEC. Deletion of either *danR* or *danI* alone also increased NET formation, though to a lesser extent than deletion of the entire DanRI system (**Figure 5E**). In summary, these results demonstrate that DanRI regulatory system dampens neutrophil activation by suppressing the ROS burst and limiting NET formation.

### The DanRI system regulates pathogenicity *in vivo*

To investigate the role of the DanRI system in UPEC pathogenicity, we intravesically infected female C57BL/6J mice with 10^7^ CFU of either the parental UPEC or Δ*danRI*, and quantified bacterial burden in the bladder at 24 and 48 hours post-infection (PI).^58^ At 24 hours PI, both strains colonized the bladder at comparable levels. However, by 48 hours PI, the Δ*danRI* strain was significantly reduced in the bladder compared to the parental UPEC strain (**Figure 6A**). These results indicate that while the DanRI system is dispensable for initial colonization, it plays a more prominent role in supporting UPEC persistence during the neutrophil response, which typically peaks around 24 hours PI.^59,60^

**Figure 6.**
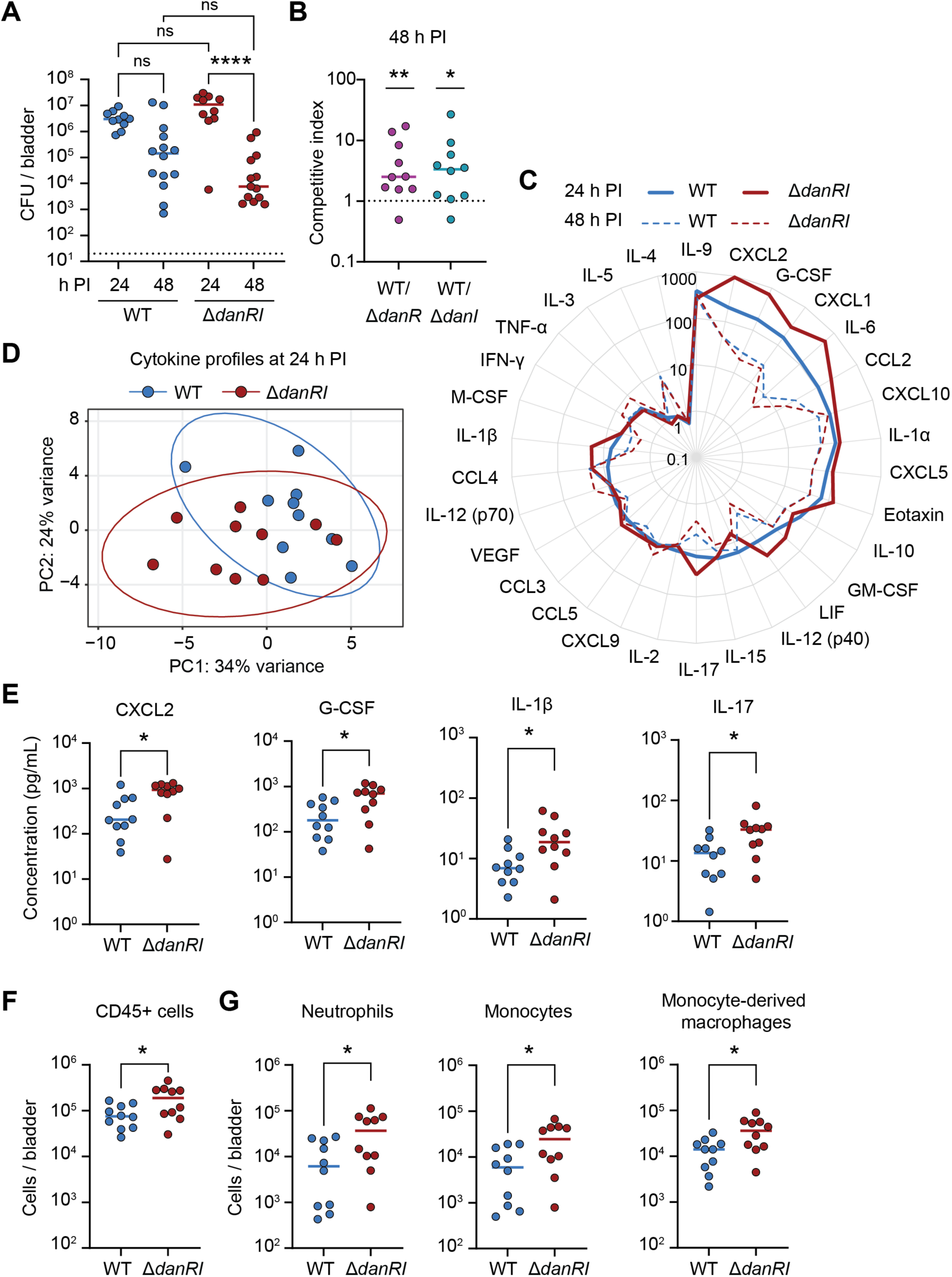
The regulatory system DanRI modulates UPEC pathogenesis. (A) Mice were infected intravesically with parental UPEC or Δ*danRI*, and CFU per bladder were determined 24 or 48 hours post-infection (median, n = 10-14 mice from 2-3 independent experiments). Significance was determined by a Kruskal-Wallis test with correction for multiple comparisons using Dunn’s test (ns - no significant difference; **** p ≤ 0.0001). (B) Mixtures (1:1) of parental UPEC and Δ*danR* or Δ*danI* were delivered intravesically into mice (median, n = 10 mice from 2 independent experiments). Bladders were harvested 48 hours post-infection, and the competitive index was calculated as CFU of parental UPEC divided by CFU of the mutant. Significance was determined with a sample Wilcoxon test to compare the median competitive index to the null hypothesis of the value of 1 (1 = no competition; * p ≤ 0.05; ** p ≤ 0.01). (C) Cytokine and chemokine expression profiles from parental UPEC and Δ*danRI-*infected bladders 24 hours (full line) and 48 hours (dashed line) post-infection (median, n = 10-14 mice from 2-3 independent experiments). (D) Principal component analysis of scaled cytokine and chemokine profiles from parental UPEC and Δ*danRI*-infected bladder at 24 hours post-infection (n = 10 mice for 2 independent experiments). An ellipse was fitted for strain sample separation. (E) Concentrations of CXCL2, G-CSF, IL-1β, and IL-17 in parental UPEC and Δ*danRI*-infected bladders 24 hours post-infection (median from 10 mice from 2 independent experiments). Significance was determined between parental UPEC and Δ*danRI* with the Mann-Whitney test corrected for the FDR (* p ≤ 0.05). (F and G) Total number of CD45+ cells (F), and of neutrophils, monocytes, and monocyte-derived macrophages (G) in parental UPEC and Δ*danRI*-infected bladders 24 hours post-infection (median from 10 mice from 2 independent experiments). Significance was determined between parental UPEC and Δ*danRI* with the unpaired t-test corrected for the FDR (* p ≤ 0.05).

To determine whether DanRI confers a fitness advantage to UPEC during infection, female mice were coinfected with equal CFU of parental UPEC and either Δ*danR* or Δ*danI.* Bladder bacterial burdens were analyzed at 48 hours PI, and competitive indexes were calculated. The parental UPEC consistently outcompeted both mutants (**Figures 6B** and **S7A**), indicating that the DanRI regulatory system enhances UPEC virulence and fitness during infection.

We next investigated whether the reduced bacterial burden observed in Δ*danRI*-infected mice is accompanied by increased inflammation. We profiled cytokines and chemokines in the bladder at 24 and 48 hours PI by Luminex. Analyte concentration peaked at 24 hours PI and declined by 48 hours (**Figure 6C**), as observed previously,^60^ which mirrored bacterial loads (**Figure 6A**). Notably, Δ*danRI*-infected bladders exhibited an increased pro-inflammatory cytokine response at 24 hours PI compared to those infected with the parental UPEC. PCA of cytokine and chemokine profiles showed notable clustering based on the infecting strain at 24 hours PI, with multivariate analysis confirming statistical significance (p = 0.0277 with PC1 and PC2) (**Figure 6D**). Among the analytes contributing most to the observed variance in principal component 1 (PC1) were GM-CSF, CXCL2, LIF, CCL2, IL-1β, G-CSF, eotaxin, and IL-17, which were all elevated in Δ*danRI*-infected bladders (**Figure S7B**). Notably, key neutrophil-recruiting and -activating factors, such as CXCL2, G-CSF, IL-1β, and IL-17, were significantly increased in Δ*danRI*-infected bladders compared to the parental UPEC infection (**Figure 6E**). These data suggest that increased inflammatory signatures at 24 hours contribute to the observed bacterial burden reduction at 48 hours.

Given the increase in expression of cytokines and chemokines that recruit innate immune cells in Δ*danRI*-infected bladders, we assessed immune cell infiltration by flow cytometry. In line with elevated cytokine expression, Δ*danRI*-infected bladders exhibited a higher number of total CD45+ cells compared to the parental UPEC-infected bladders at 24 hours PI (**Figure 6F**). Within the total CD45+ immune cell population, infiltrating neutrophils, monocytes, and monocyte-derived macrophages were significantly increased in Δ*danRI*-infected bladders (**Figure 6G**), while resident macrophage numbers remained unchanged (**Figure S7C**). Using RFP-expressing bacteria, we evaluated bacterial uptake by immune cells. No significant differences in bacterial uptake were observed by neutrophils or monocyte-derived macrophages (**Figure S7D**), consistent with our *in vitro* phagocytosis experiments (**Figure S6A**). Interestingly, monocytes exhibited significantly greater uptake of the Δ*danRI* mutant (**Figure S7D**), suggesting that in the absence of DanRI, UPEC becomes more vulnerable not only to neutrophil-mediated clearance but also to monocyte-mediated uptake.

Taken together, these findings suggest that the DanRI system attenuates early immune cell recruitment and activation in the bladder, giving UPEC a survival advantage. In its absence, the observed increase in immune cell recruitment and neutrophil responses may contribute to enhanced bacterial clearance, suggesting a potential role for DanRI in modulating host-pathogen interactions during UTI.

## Discussion

UPEC must adapt to the dynamic environment of the urinary tract, including the infiltration of neutrophils and the deployment of NETs. Here, we investigated how UPEC responds to NETs and demonstrated that, unlike commensal *E. coli*, UPEC can survive NET-mediated killing. This resilience likely stems from evolved mechanisms that specifically counteract neutrophil functions.^61^ Indeed, several known UPEC virulence factors suppress neutrophil functions,^26,27^, for example, TcpC can inhibit the release of NETs.^28^ These adaptations likely contribute to its success as a uropathogen.

We observed that, in contrast to other pathotypes, UPEC initiates a robust transcriptional response upon exposure to NETs. This includes the upregulation of pathways involved in overcoming metal limitation and oxidative stress, as well as metabolic reprogramming to conserve energy. Notably, UPEC upregulated two siderophore systems, yersiniabactin and salmochelin, despite their high metabolic cost.^62^ These systems likely counteract abundant NET-associated host proteins, such as lactoferrin and calprotectin, which sequester essential metals.^19^ This molecular ‘tug of war’ over metal availability between UPEC and the host is critical for UPEC survival during infection.

Transcriptomic analysis revealed a previously uncharacterized two-gene regulatory system, which we named DanRI. We propose that the regulator, DanR, modulates transcription by binding to DNA via its LytTR-like domain, while DanI inhibits DanR transcriptional activity. This regulatory configuration, in which one protein is a transcriptional regulator and the other is an inhibitor, is reminiscent of the HdrRM and BrsRM systems of *Streptococcus mutans,*^63,64^ recently classified as LytTR Regulatory Systems (LRS).^65^ These systems are notable for their unique modes of regulation, which differ from classical bacterial regulatory pathways. Unlike most LRS systems, where gene expression is typically regulated by the LytTR regulator itself, DanRI appears to lack such autoregulatory feedback. Additionally, while LRS systems are distinct from canonical two-component systems composed of a histidine kinase sensor and a response regulator,^29^ DanRI also lacks the hallmark sensor-regulator pairing. Other two-component response regulators with LytTR domains, such as AgrA in *Staphylococcus aureus*^35^ and VirR in *Clostridium perfringens*,^66^ use additional domains to receive input from sensor kinases. Such domains are absent in both DanR and LRS-type regulators. Interestingly, LytTR domains are predominantly associated with Gram-positive bacteria,^67^ suggesting that the discovery of DanR expands our understanding of the diversity of transcriptional regulators.

Is DanI a sensor in the DanRI system? DanI is predicted to contain a four-cysteine zinc finger domain, which, although classically associated with DNA binding, is increasingly recognized for mediating protein-protein and lipid interactions.^68^ The zinc finger of DanI is homologous to that of DksA,^41^ a redox-sensitive protein that responds to reactive oxygen and nitrogen species via conformational changes.^44^ Given that UPEC lacking DanRI triggers an increased ROS response in neutrophils, DanI may function as a redox-sensitive sensor that detects ROS via its cysteine residues and regulates DanR activity accordingly, ultimately modulating neutrophil activation.

The DanRI system is common among UPEC strains but rare in other *E. coli* pathotypes, with only distant homologs identified in a few other bacterial species. Notably, 29% of UPEC strains encode DanRI distributed across distinct clonal lineages. Like DanRI, other regulatory systems, such as KguS/R,^69^ OrhK/R,^70^ GrpP/Q^71^, and RcrARB,^72^ are also predominantly associated with UPEC, yet not universally present. This diversity might reflect the diverse host niches occupied by UPEC, where strain-specific regulatory systems provide adaptive advantage to environmental challenges.^29^ The presence of DanRI may also serve as a genetic marker distinguishing UPEC from commensal *E. coli* strains, addressing a current challenge in the field.

Functionally, DanRI modulates the host immune response by attenuating the ROS burst, a critical trigger for NET formation. Neutrophil-derived ROS can affect microbes extracellularly, be delivered into phagolysosomes, or trigger intracellular signals leading to NET release.^73^ Notably, ROS generated by NADPH oxidase have recently been implicated in UPEC clearance,^13^ which aligns with our findings that pharmacological inhibition of this enzyme dampens the oxidative response to UPEC. Importantly, infections with DanRI-deficient UPEC result in increased NET formation, consistent with their impaired ability to suppress ROS production.

*In vivo* infections revealed a distinct role for the DanRI regulatory system in modulating the innate immune response during bladder infection. Temporal infection dynamics showed that DanRI is not required for initial bacterial colonization but becomes critical as the host immune response intensifies. At 24 hours PI, we observed increased cytokines associated with neutrophil recruitment and activation, such as G-CSF, which promotes neutrophil granulopoiesis,^10^ and CXCL2, a chemokine that mediates neutrophil recruitment.^74^ Increased cytokine expression in Δ*danRI*-infected mice was correlated with greater myeloid cell infiltration, including neutrophils, at 24 hours PI, which led to reduced bacterial burden at 48 hours PI. Indeed, an early, robust immune response is critical to prevent the development of chronic UTI.^60,75^ Additionally, IL-17, a cytokine necessary for UTI resolution,^60^ was significantly increased in the Δ*danRI* infection. IL-17 is a multifunctional mediator that regulates inflammatory pathways, including chemokine production,^76^ and drives neutrophil recruitment and activation.^77^ Its elevated expression suggests that γδ T cells, Th17 cells, or ILC3s contribute to a feed-forward inflammatory loop sustaining neutrophil infiltration and/or activity.^60,78^ These findings underscore the key role of neutrophils as a front-line defense in the bladder, and highlight the role of DanRI in modulating early host responses and shaping infection trajectories favorable to UPEC persistence.

Mechanistically, how DanRI exerts its immunomodulatory effect remains to be elucidated. DanRI regulates multiple pathways, including nutrient transport, metabolism, stress responses, S-fimbriae, and flagellum biosynthesis. For instance, *hdeD,* which is upregulated in Δ*danI*, can suppress flagellar expression,^79^ while *ymgA,* which is downregulated in Δ*danR,* can activate the RcsCDB two-component system during stress.^80^ These examples highlight the complexity of the DanRI regulatory network, which likely operates directly and indirectly across multiple pathways within the bacteria.

Targeting the DanRI system in UPEC could potentiate current treatments by impairing the ability of UPEC to evade host defenses and reduce persistence. Given the rise in antibiotic resistance and the persistence of UPEC infections, manipulating regulatory networks that modulate host-pathogen interactions offers a promising direction for future research. The presence of NET-associated proteins in urine^18^ further reinforces the relevance of this immune response in UTI and highlights the need to explore how NETs can be harnessed for better bacterial clearance.

In summary, we identified DanRI, a novel regulatory system in UPEC that modulates neutrophil-mediated host defenses to promote bacterial persistence. We propose a model in which UPEC, upon exposure to NETs and NET-derived nucleosomes in the urinary tract, upregulates the DanRI to rapidly initiate a protective response against infiltrating neutrophils. Through the regulation of essential pathways, including motility, nutrient transport, metabolism, and stress responses, DanRI suppresses inflammatory cytokine expression and dampens neutrophil activation, thereby enabling immune evasion and prolonged survival in the host. Altogether, DanRI represents a previously unrecognized strategy through which UPEC fine-tunes its response to innate immune pressure, underscoring the sophisticated nature of bacterial adaptation during infection.

## Methods

### Ethics statement

Experiments using human samples were conducted in accordance with the Declaration of Helsinki. Blood samples were collected from anonymous donors with consent and approval from the Charité Hospital ethics committee in Berlin, Germany (EA1/0104/06). Mouse infection experiments were conducted in accordance with the approval of the protocol number APAFIS #34290 by SC3 - CEEA34 – Université de Paris Cité, at the Institute Cochin in application of the European Directive 2010/63 EU.

### Bacterial strains and culturing conditions

All bacterial strains (**Table S1**) were stocked in Luria-Bertani (LB) with 10% glycerol at -80°C. Bacterial cultures were streaked from stock on LB agar plates and, unless stated otherwise, one colony from a fresh plate was inoculated in 5 ml of LB and grown overnight at 37°C, shaking. The overnight suspension was diluted 1:100 into fresh media and grown for 2 hours at 37°C to reach the logarithmic phase. Cultures were centrifuged for 5 minutes at 2,000 x g and washed twice with RPMI 1640 (pH 7.4, with L-glutamine, without phenol red; Sigma Aldrich) supplemented with 10 mM HEPES (Gibco). For experiments with neutrophils and NETs, the same medium was used with the addition of 0.5% human serum albumin (Bio&SELL). Bacterial concentrations were estimated by measuring the optical density at 600 nm (OD_600_). When needed, kanamycin was used at 50 µg/mL, chloramphenicol at 34 µg/mL, and carbenicillin at 100 µg/mL, unless stated otherwise.

### Human neutrophil isolation

The peripheral blood of healthy adult donors was collected in EDTA-containing tubes. The blood was layered on an equal volume of Histopaque 1119 (Sigma-Aldrich) and centrifuged for 20 minutes at 800 x g with acceleration 7 and deceleration 3 - 5. The pink layer containing neutrophils was collected and washed with Dulbecco’s phosphate-buffered saline (DPBS, Gibco) supplemented with 0.1% human serum albumin for 10 minutes at 300 x g. The resulting pellet was resuspended in 2 mL DPBS with 0.1% human serum albumin and layered on a discontinuous Percoll gradient (Cytiva) diluted in DPBS with decreasing densities starting at 85% on the bottom, then 80%, 75%, 70%, and 65% in 2 mL layers in a 15 mL tube. After 20 minutes of centrifugation at 800 x g with acceleration 7 and deceleration 3 - 5, the neutrophils were collected from the interface between 65% and 75% of the gradient. Neutrophils were washed again with DPBS with 0.1% human serum albumin and counted with CASY cell counter (OMNI Life Science). Purified neutrophils were used immediately after isolation.

### NETs killing assay

In a flat-bottom 96-well plate, 5 x 10^5^ neutrophils in 50 µL were seeded and stimulated with 100 nM phorbol 12-myristate 13-acetate (PMA, Sigma-Aldrich) or 30 µM nigericin (Sigma-Aldrich), as previously described.^81^ The plate was incubated for 4 hours at 37°C. Neutrophils were incubated with 10 µg/mL of cytochalasin B (Thermo Fisher Scientific) for 15 minutes at 37°C before adding bacteria. Bacterial cultures were used at 5 x 10^6^ in 50 µL for a multiplicity of infection of 10. Bacteria were incubated with 10 µg/mL of rifampicin (ITW Reagents) for 15 minutes at 37°C before the addition to the NETs. Bacterial cultures were added to wells, the plate was centrifuged for 5 minutes at 800 x g, and incubated for 1 hour at 37°C. As a control, bacterial cultures were incubated in a medium with the addition of PMA or nigericin. To collect the bacteria, 20 U/mL of micrococcal nuclease (Sigma or TakaraBio) was added for 10 minutes, and the wells were resuspended. Subsequently, the samples were serially diluted 1:10 in DPBS and plated on LBA plates. The plates were incubated at 37°C overnight and CFU were counted. All conditions were performed in technical duplicates. Bacterial killing was calculated as a percentage of CFU obtained from bacteria on NETs divided by bacteria in the medium with stimulus.

### RNA isolation and sample preparation

In a 12-well plate, 500 µL of 5 x 10^7^ bacteria were added to wells with 500 µL medium RPMI with 0.5% human serum albumin with stimuli or with NETs. To generate NETs, 5 x 10^6^ neutrophils were stimulated with 100 nM PMA for 4 hours at 37°C, and the NETs were either left intact or digested with 10 U/mL of DNase I (New England BioLabs). Conditions were performed in technical duplicates that were combined during RNA isolation. The plate was centrifuged at 800 x g for 5 minutes and incubated for 1 hour at 37°C. To collect bacteria, NETs were processed with 10 U/mL DNase I (New England BioLabs) for 10 minutes, resuspended, and scraped. All samples were collected in a tube, centrifuged at 12,000 x g for 5 minutes, and the pellet was resuspended in RNA later stabilization solution (Invitrogen) for 1 hour at room temperature. To isolate the RNA, the tubes were centrifuged, and the pellet was resuspended in TE buffer (10 mM Tris and 1 mM EDTA in dH_2_0) with 50 µg/mL of lysozyme (Thermo Fisher Scientific) and 2 mg/mL of proteinase K (Qiagen). The tubes were incubated for 10 minutes at room temperature with 700 rpm shaking to lyse the bacteria. RNA was isolated with the RNeasy mini kit (Qiagen) according to the manufacturer instructions. The samples were additionally processed with on-column DNase I digestion (Qiagen) to remove contaminating DNA. RNA concentration was measured on a Nanodrop 2000 (Thermo Fisher Scientific) or using Broad Range Qubit RNA assay (Thermo Fisher Scientific). RNA was stored at -80°C.

### RNA sequencing and analysis

Total RNA was quality controlled using an Agilent RNA 6000 Nano kit on the 2100 Bioanalyzer system (Agilent). 1 µg of total RNA with an RNA integrity number above 8 was processed using the Ribo-Zero Plus rRNA Depletion kit (Illumina) to remove eukaryotic and prokaryotic rRNA. RNA sequencing of NETs and UPEC (5 biological replicates) was done by Eurofins (INVIEW Transcriptome Bacteria), and the sequencing of UPEC mutants (3 biological replicates) by Novogene. Companies conducted the library preparation and Illumina sequencing with a 150 bp paired-end read length. Approximately 10 million paired reads were obtained per sample and controlled for quality using FastQC and MultiQC. Bowtie2 was used to align reads to the reference genome of *E. coli* UTI89 (NC_007946.1^37^) using default settings.^82^ Reads mapping to each gene in the assembly (GCF_000013265.1) were counted using featureCounts.^83^ Counts were normalized with SizeFactors, and DESeq2 was used for differential gene expression analysis with the unstimulated UPEC or the wildtype UPEC set as the reference level.^84^ The threshold for significance was set to adjusted p-value < 0.05 and log2 fold change threshold > 0.5. Log2 fold changes were shrunk using the lfcShrink with the type apeglm model.^85^ Visualization was performed in R Studio software and included hierarchical clustering, principal component analysis, heatmaps, and volcano plots to explore sample correlations and expression changes.

### Gene ontology enrichment analysis

Gene ontology terms from the differentially expressed genes were analyzed for over-representation with enrichGO in RStudio using the *E. coli* K12 (org.EcK12.eg.db) database.^34^ The analysis focused on Biological Processes. The results were adjusted for false discovery rate using the Benjamini–Hochberg method with significant pathways identified at a p-value cut-off of 0.05. The enriched terms were further processed with Simplify, with a cut-off set at 0.6 to remove redundant results.

### RT-qPCR

High-Capacity cDNA Reverse Transcription Kit (Applied Biosystems) was used to reverse transcribe 100 ng of total RNA into cDNA using random primers according to the manufacturer instructions. Quantitative PCR was performed in MicroAmp Fast Optical 96-Well Reaction Plate (Thermo Fisher Scientific) using the Fast SYBR Green Master Mix (Thermo Fisher Scientific), 1 µL of cDNA, and 500 nM of forward and reverse primers for genes of interest (**Table S2**). Each condition was set up in triplicate. QPCR was run using the fast program settings in the QuantStudio 3 Real-Time PCR System (Applied Biosystems). Negative controls included no template and no reverse transcriptase. Results were analyzed using the Quant Studio Design and Analysis Software v1.5.2. The cycle of the threshold value (C_t_) was obtained for each condition, normalized to the expression of housekeeping gene β-subunit of gyrase (*gyrB*), and expressed as relative fold change compared to the control using the 2^-ΔΔCt^ method.

### Blast analysis, sequence alignment, and phylogeny

The sequences were analyzed with the TBLASTX tool against *E. coli* genomes available in the PubMLST database.^45^ Results were processed in R Studio, with genomes considered positive for sequences if they showed a hit with >50% identity. Protein sequences were copied from NCBI^39^ and aligned using the multiple sequence alignment program Clustal Omega, clustered by neighbor-joining. The analysis and the generation of the phylogenetic tree were done through the EMBL European Bioinformatics Institute services.^86^

### Construction of deletion mutants

The α-Red recombinase site-specific insertion protocol was used to generate mutant strains and was adapted from previous studies.^52,53^ Plasmid pKD4 was used to amplify a kanamycin resistance cassette and plasmid pKD3 for a chloramphenicol cassette (**Table S3**). PCR was performed using the Q5 High-Fidelity DNA Polymerase (New England Biolabs) and primers P1 and P2 (**Table S2**) with 50 bp overhangs complementary to the 5’ and 3’ ends of the insertion site. PCR products were digested with 20 U of DpnI (New England Biolabs), analyzed on 1% agarose gel, purified using DNA Clean & Concentrator (Zymo Research), and quantified. UPEC with plasmid pKM208 (**Table S3**) was grown in LB with carbenicillin at 30°C with shaking, induced with 1 mM isopropyl β-d-1-thiogalactopyranoside (Carl Roth) at an OD_600_ of 0.01, and further grown until OD_600_ of 0.1. Subsequently, the culture was incubated at 42°C for 15 minutes and on ice for 10 minutes. Cultures were centrifuged and resuspended in decreasing volumes of ice-cold 20% glycerol in 1 mM MOPS buffer to prepare competent cells, which were frozen at -80°C. For electroporation, cells were thawed, mixed with 300 - 500 ng of DNA, and shocked using a MicroPulser Electroporator (Bio-Rad). Post-shock, cells were incubated in 1 mL of S.O.C medium (New England Biolabs) for a minimum of 90 minutes at 37°C, spread on LBA plates with 25 µg/mL of kanamycin or chloramphenicol, and incubated overnight. To remove the antibiotic cassette, plasmid pCP20 (**Table S3**) was electroporated, and cultures were incubated in S.O.C medium at 30°C for 1 hour, spread on carbenicillin LBA plates, and incubated overnight at 30°C. The culture was grown at 43°C to induce FLP recombinase and eliminate pCP20. Verification of cassette removal and correct site-specific deletions was performed via colony PCR and sequencing.

### Cloning of plasmids

All cloning of plasmids (**Table S3**) was done using In-Fusion Snap Assembly Master Mix (Takara Bio). PCR to amplify gene and promoter sequences was done using the Q5 High-Fidelity DNA Polymerase (New England Biolabs) and primers for specific genomic locations (**Table S2**). Plasmid pTrc99a (**Table S3**) was used to generate complementation strains, in which the sequence of the whole region or each gene was inserted into the multiple cloning site under the control of the *trc* promoter and *lac* operator, and the plasmids were transformed into the respective mutants. Plasmid pGEN-luxCDABE (**Table S3**) was used to create a reporter system to measure promoter activity. The plasmid was digested with PmeI and SnaBI (New England Biolabs) to remove the constitutive promoter sequence, and PCR-amplified promoters of genes were inserted into the site. Promoters were predicted in intergenic regions larger than 40 bp and defined from 500 bp upstream to 150 bp downstream of the transcription start site, as previously described.^87^ All constructs were verified with Sanger sequencing.

### Bacterial growth profiles

Overnight bacterial cultures were adjusted to an OD_600_ of 0.05 in LB or pooled human urine (Innovative Research), which was filter sterilized. 100 µL of culture was added to a flat-bottom 96-well plate in technical triplicates. As a control, wells with just media were used. Plates were placed into an H1 plate reader (BioTek) preheated to 37°C, and absorbance at 600 nm was measured every 10 minutes for 12 hours at 37°C with continuous linear shaking at 493 cycles per minute.

### Luciferase reporter assay

In 96-well black plates with optical bottom (Thermo Fisher Scientific), 50 µL of 2.5 x 10^6^ bacteria with pGEN plasmids (Table S1 and Table S2) was added with 50 µL of stimuli or left unstimulated. Stimuli included: NETs from 2.5 x 10^5^ neutrophils stimulated with 100 nM PMA for 4 hours, NET-purified nucleosomes obtained through in-column purification per the manufacturer’s instructions (Sartorius ultrafiltration products); recombinant histones: H1^0^, H2A, H2B, H3.1, and H4 (New England BioLabs); recombinant proteins: myeloperoxidase, neutrophil elastase, calprotectin, cathelicidin, cathepsin G, and α-defensins (Abbexa); and 100 ng of human genomic DNA (Sigma-Aldrich). All proteins were used at 10 nM, and all conditions were in RPMI with 0.5% human serum albumin in technical duplicates. Plates were incubated for 1 hour at 37°C, after which the luminescence signal was measured in a Victor X Light luminometer (Perkin Elmer). Fold changes were obtained by normalizing the luminescence units of stimulated to unstimulated bacteria from one experiment.

### Neutrophil phagocytosis assay

Neutrophils in 50 µL of RPMI were seeded into 1 mL 96-well plates at a density of 10^5^ cells per well and treated with 10 µg/mL of cytochalasin B or were left untreated for 15 minutes at 37°C. Bacterial cultures were opsonized in RPMI with 10% heat-inactivated human serum (Sigma-Aldrich) for 20 minutes at 37°C with shaking and were added in 50 µL to neutrophils at multiplicity of infection of 10. All conditions were performed in technical duplicates. Controls included bacterial incubation in media with the addition of cytochalasin B and in media alone. The plate was centrifuged at 2,000 rpm for 3 minutes and incubated for 30 minutes at 37°C with slow shaking. After incubation, the wells were resuspended, and 900 µL of ice-cold H_2_0 at pH 11 was added for 10 minutes. The suspension was serially diluted 1:10 in DPBS, plated on LBA plates, and incubated overnight. The CFU were counted the next day. Bacterial survival was calculated as a percentage of CFU obtained from bacteria with neutrophils divided by bacteria with neutrophils treated with cytochalasin B.

### ROS chemiluminescence assay

Bacterial cultures were incubated in RPMI with 10% heat-inactivated human serum for 20 minutes at 37°C with shaking. Neutrophils were seeded in a flat-bottom white 96-well plate at 10^5^ cells per well in 50 µL and incubated for 15 minutes at 37°C to allow them to settle. Diphenyleneiodonium chloride (DPI, Calbiochem) was used at 1 µM. 1.2 U/mL of horseradish peroxidase (Thermo Fisher Scientific) and 50 µM luminol (Sigma-Aldrich) were added to the wells, and the plate was incubated for 5 minutes at 37°C. Subsequently, 50 µL of 10^6^ opsonized bacteria were added to the wells. As a negative control, neutrophils were left unstimulated, and as a positive control, neutrophils were stimulated with 100 nM PMA. The plate was immediately added to the Victor Light luminescence counter (Perkin Elmer), and luminescence was measured every 2 minutes for 3 hours at 37°C. Conditions were performed in technical triplicates, and the area under the curve of the averages was calculated for each condition. Values of one experiment were normalized by minimum-maximum scaling.

### Immunofluorescence staining and microscopy

Neutrophils were seeded in 8-well µ-slide dish (ibidi) at a density of 10^5^ in 100 µL per well in medium, 100 nM PMA, or 10^6^ bacteria for multiplicity of infection of 10 in 100 µL. The dish was centrifuged at 2,000 rpm for 3 minutes and incubated for 4 hours at 37°C. The conditions were performed in technical duplicates. The wells were fixed with 2% paraformaldehyde (Electron Microscopy Sciences) for 20 minutes at room temperature, washed with DPBS, blocked in 10% normal goat serum (Sigma-Aldrich) and 1% bovine serum albumin in DPBS for 30 minutes. The samples were stained with 2 µg/mL of primary mouse monoclonal antibody 3D9recognizing NETs (in-house production)^57^ for 1 hour, followed by several washes in DPBS and incubation with 2 µg/mL of secondary antibody goat anti-mouse labelled with Alexa Fluor 647 (RRID: AB_2535804, Thermo Fisher Scientific) for 1 hour. Finally, the wells were washed twice with DPBS, stained with 0.5 µg/mL of 4’,6-diamidino-2-phenylindole (DAPI, Invitrogen) for 5 minutes, and washed with DPBS. Images were acquired with a 20X objective on an EVOS M7000 Imaging System (Invitrogen). Fiji software was used to process the images.^88^

### NETs ELISA

Neutrophils were seeded into a 12-well plate at 10^6^ per well and were left unstimulated, or were stimulated with 100 nM PMA, as a positive control. Bacteria at a multiplicity of infection of 10 were added to neutrophils, the plate was centrifuged at 2,000 rpm for 3 minutes, and incubated for 4 hours at 37°C. Conditions were performed in technical duplicates. 20 U/mL of micrococcal nuclease was added for 10 minutes, and the wells were scraped and resuspended. The suspension was spun down at 300 x g for 5 minutes to remove cell debris, and the supernatant was stored at -20°C. ELISA was performed using the SimpleStep ELISA Custom ELISA Kit (Abcam) following the manufacturer’s protocols. The antibody 3D9 (in-house production)^57^ was conjugated with the capture peptide, and a monoclonal MPO antibody (RRID: AB_934783, Thermo Fisher Scientific) was conjugated with HRP. Samples were thawed, and 50 µL was added to the microplate strips in duplicates as undiluted and diluted 1:2 in sample dilution buffer for secreted proteins from the kit. Both antibodies were added to the wells at a final concentration of 200 ng/mL of capture-NETs 3D9 antibody and 100 ng/mL of HRP-MPO. The plate was incubated for 1 hour with shaking at 600 rpm and washed 5 times with the wash buffer. A detection reagent was added immediately afterwards, the plate was incubated for 15 minutes with shaking at 600 rpm, and the stop solution was added. The absorbance at 450 nm was measured in the H1 plate reader (BioTek). The data were adjusted for background signals from wells with only antibodies.

### Mouse infections

Six- to seven-week-old female C57BL/6J mice (Charles River Laboratories, France) were used for all infection experiments. For single-strain infections, mice were infected with either the parental UPEC strain UTI89 attB::Km-marsRFP^59^ or the mutant UTI89Δ*danRI*::Km-marsRFP. For competitive index infections, mice were infected with the parental UTI89 attB::bla-GFP and either UTI89Δ*danR*::Km-marsRFP or UTI89Δ*danI*::Km-marsRFP at equal concentrations. Bacterial cultures were initiated by inoculating single colonies into 10 mL LB supplemented with appropriate antibiotics, followed by overnight incubation at 37 °C without shaking. The next day, bacteria were pelleted by centrifugation and resuspended in PBS for infection. The accuracy of the inoculum was confirmed by serial dilution and plating on LB agar. The mixed inoculum (50 µL total, 10⁷ CFU) was prepared by combining equal CFU of each strain. Mice were anesthetized via intraperitoneal injection of 100 mg/kg ketamine and 5 mg/mL xylazine, catheterized transurethrally, and infected with 50 µL of bacterial suspension in PBS with 10⁷ CFU total. At 24 or 48 hours post-infection, mice were euthanized by cervical dislocation. Bladders were aseptically removed, placed into 1 mL PBS, and homogenized using a PreCellys bead homogenizer (Bertin Technologies). Homogenates were serially diluted 1:10 in PBS and plated on LB agar containing appropriate antibiotics. Plates were incubated overnight at 37 °C, and CFU were enumerated the following day. The detection limit was 20 bacteria per bladder. For CI experiments, the competitive index was calculated for each bladder as: CI = (mutant CFU / parental CFU).

### Luminex xMAP analysis

Homogenized bladders from mouse infections were centrifuged at 17 x g for 5 minutes at 4°C. The supernatant was transferred to a V-bottom low protein binding plate and immediately frozen at -20°C for future analysis. Cytokine and chemokine concentrations were quantified using the MILLIPLEX MAP mouse premixed magnetic bead panel for 32 analytes (Merck Milipore), according to the manufacturer’s recommendations. To minimize batch effects, all samples were measured concurrently. Analytes IL-13 and IL-7 were excluded from the analysis because they had concentrations below the minimal detectable concentration plus 2 standard deviations, as per the manufacturer’s recommendations.

### Flow cytometry

Bladders from mouse infections were collected, minced with scissors, and digested with 0.34 U/mL liberase in PBS for 1 hour at 37°C.^58^ To stop the digestion, PBS supplemented with 2% fetal bovine serum and 0.2 mM EDTA (FACS buffer) was added to each sample. Cell suspensions were passed through a 100 μm filter (Miltenyi Biotec), and the filters were washed with FACS buffer. The tubes were centrifuged at 1,200 rpm for 7 minutes at 4°C, pellets were resuspended in FACS buffer and transferred to a 96-well plate. Plates were centrifuged at 1,200 rpm for 5 minutes at 4°C, the pellet was resuspended in 100 µL of FACS buffer and 1:100 of FC block (anti-CD16, CD32) and incubated for 15 minutes at 4°C. 100 µL of antibody cocktail (antibodies diluted 1:100, **Table S4**) in brilliant stain buffer (BD Bioscience) was added to each well, and the plate was incubated for 30 minutes at 4°C, protected from light. The plate was centrifuged at 1,200 rpm for 5 minutes at 4°C, resuspended in 4% paraformaldehyde and incubated for another 10 minutes. Finally, the plate was centrifuged at 1,200 rpm for 5 minutes at 4°C, and the samples were resuspended in FACS buffer. Total cell counts were determined with AccuCheck counting beads as previously described.^59^ Flow cytometry was performed on an LSR Fortessa and data were analyzed using FlowJo (TreeStar) software.

### Statistical analysis

Details of sample size and statistical methods used for each experiment are indicated in the figure legends. Statistical analysis was performed in GraphPad 10.1.2 or RStudio software. When mentioned, additional multiple testing correction was performed with the false discovery rate method of Benjamini and Hochberg (q = 5%). Statistical significance was set for all data at a p-value or q-value ≤ 0.05.

## Materials availability

All unique/stable reagents generated in this study are available from the lead contact with a completed material transfer agreement.

## Data and code availability

Analyzed RNA-sequencing results are available as supplementary data to this study. RNA-sequencing raw reads generated for this study have been deposited in the European Nucleotide Archive (ENA) at EMBL-EBI under accession number PRJEB89779. R scripts used for analysis and generating graphs are available at github.com/doracerina/UPEC-NETs. This study did not generate any original code. Any additional information required to reanalyze the data reported in this study is available from the lead contact upon request.

## Acknowledgments

We thank all the past and present members of the Zychlinsky lab and the Ingersoll lab for insightful discussions and suggestions. We thank Volker Brinkmann, Christian Goosmann, and Robert Hurwitz for support in this study. We thank CJ Harbort for the help with R analysis. We thank all the blood donors for their contributions. This work was funded by the Max Planck Society (A.Z.) and the *Agence Nationale de la Recherché* (French National Research Agency) ANR-19-CE15-0015 (M.A.I.).

## Author contributions

Conceptualization by D.Č., G.M., A.Z., and M.A.I.; Investigation and analysis by D.Č., M.R., and C.H.O.; Writing of the original draft by D.Č.; Reviewing and editing by D.Č., M.R., A.Z., and M.A.I.; Funding by A.Z., and M.A.I.; Supervision by G.M., A.Z., and M.A.I.

## Supplemental information

### Supplemental figures

**Figure S1.**
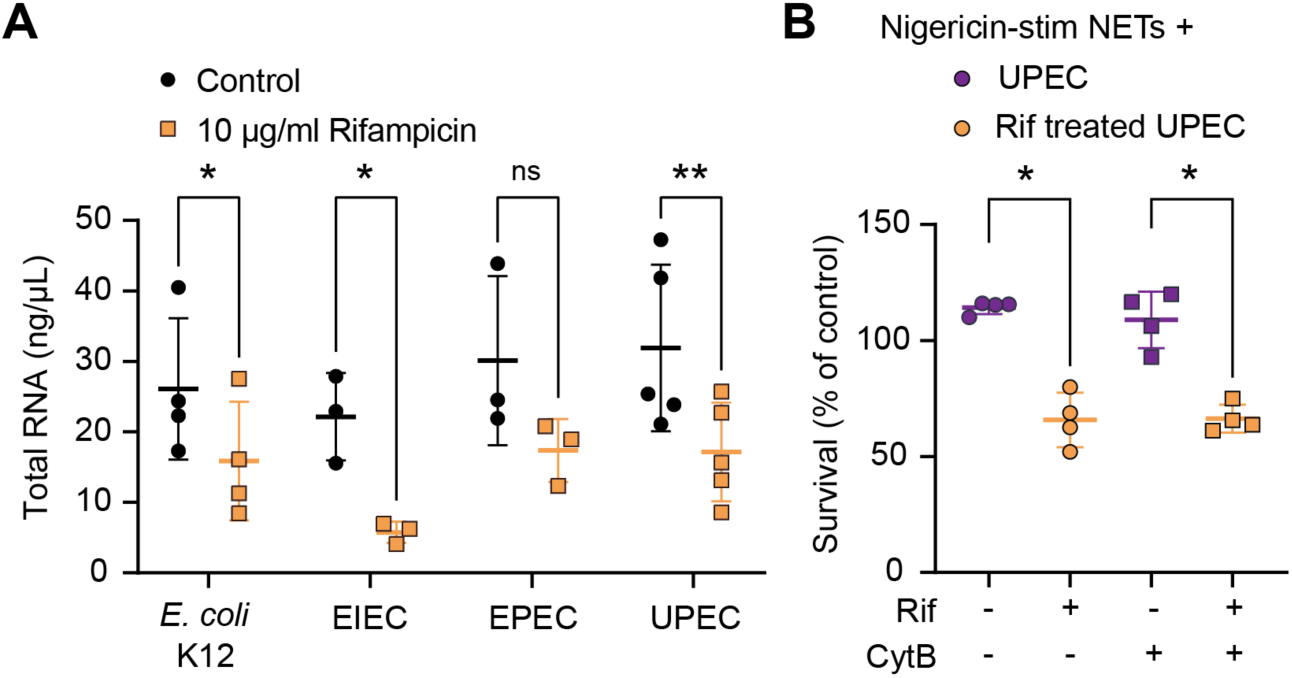
Rifampicin decreases RNA and impacts UPEC’s survival on NETs. (A) Total RNA was isolated from bacteria incubated for 1 hour with or without 10 µg/mL of rifampicin (mean ± SD, n = 3-4). Significance was determined with a paired t-test (ns - no significant difference, * p ≤ 0.05; ** p ≤ 0.01). (B) Nigericin-stimulated NETs were treated with or without 10 µg/mL cytochalasin B to inhibit phagocytosis. UPEC was treated with or without 10 µg/mL of rifampicin and was added to NETs or medium with stimuli for 1 hour at an MOI of 10. Survival is expressed as a percentage of CFU of UPEC with NETs normalized to the appropriate controls of UPEC in medium with stimuli (mean ± SD, n = 4). Significance was determined with a Mann-Whitney test between control and stimulated condition (* p ≤ 0.05).

**Figure S2.**
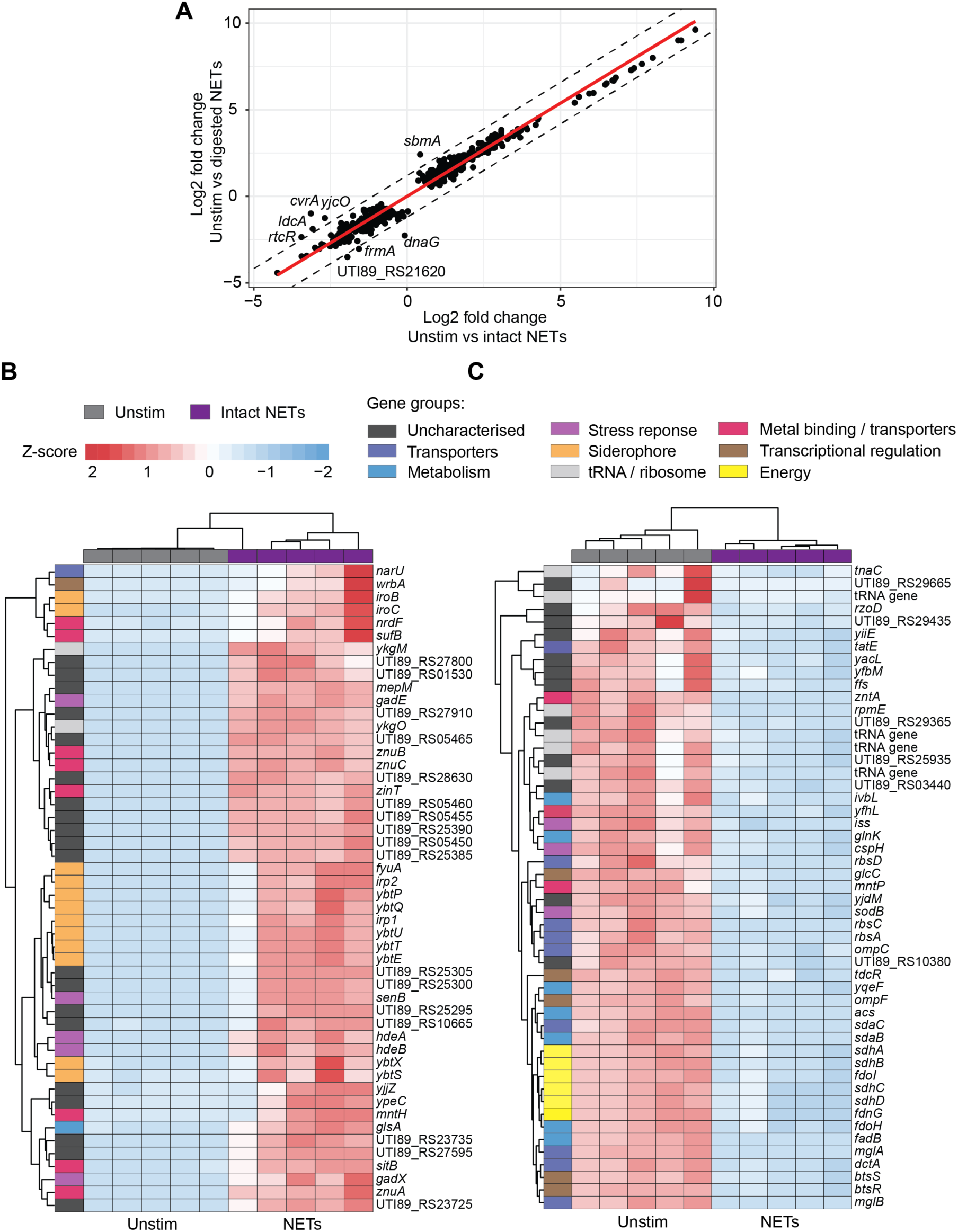
NETs interaction changes gene expression in UPEC. (A) Correlation plot of log2 fold changes of differentially expressed genes in UPEC after incubation with intact NETs (x-axis) and DNase-digested NETs (y-axis). The red line represents the linear regression fit, while the dotted lines indicate a 1.2 threshold distance in log2 fold changes. (B and C) Heatmap displaying Z scores of top 50 upregulated (B) and downregulated (C) differently expressed genes in UPEC exposed to intact NETs compared to the unstimulated condition. The genes were clustered based on their normalized gene expression and were matched to 9 functional categories based on literature searches: unknown/other, transporters, metabolism, stress response, siderophore, tRNA/ribosome, metal binding/transporters, transcriptional regulation, and energy production.

**Figure S3.**
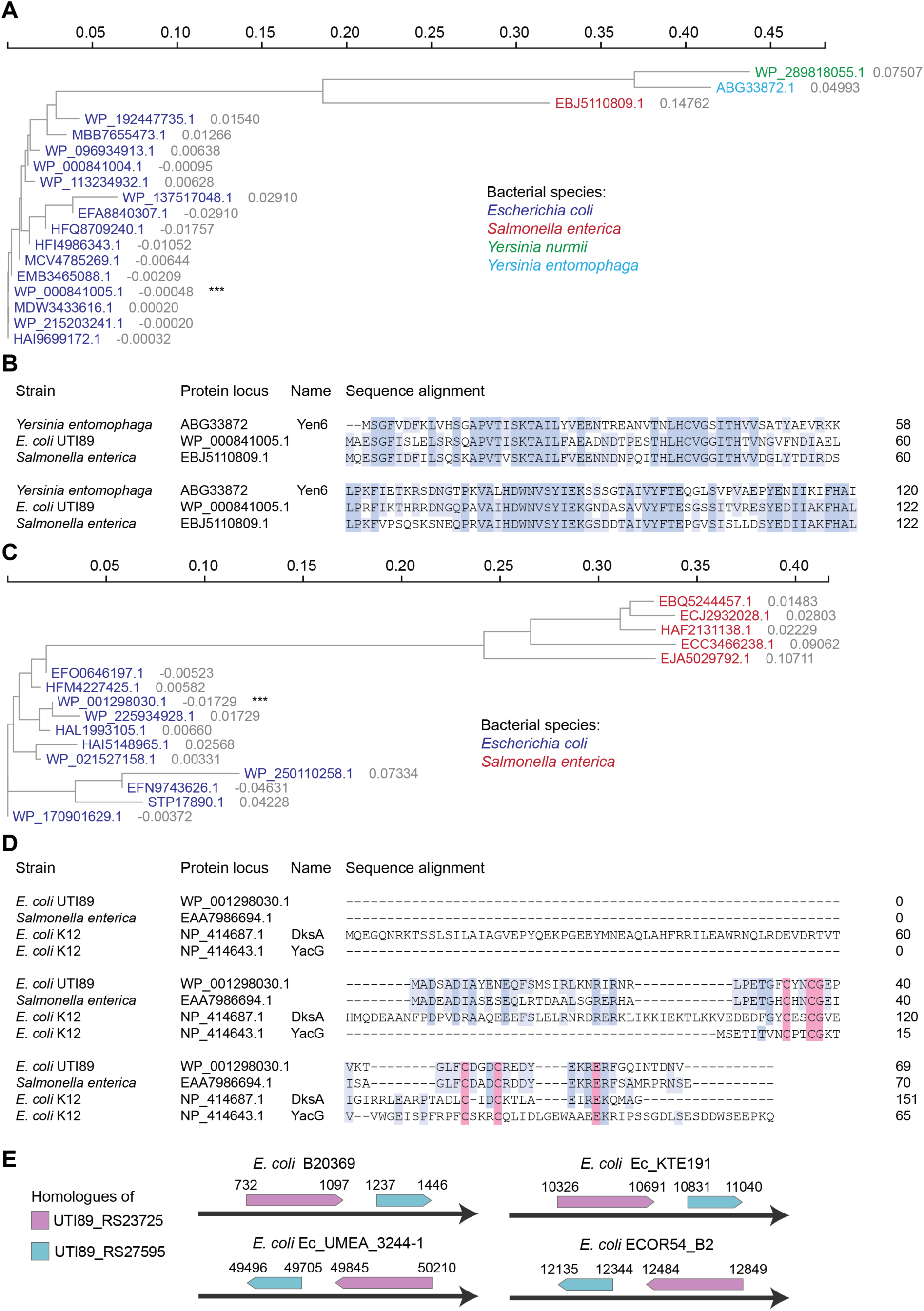
Homologues are present in *E. coli* and other bacterial species. (A and C) Phylogenetic tree of proteins aligned using the Clustal Omega and clustered by the neighbor-joining method. The distance between the sequences is shown in grey, and the protein IDs are color-coded based on their bacterial species. Protein sequences with more than 50% identity to the protein from *E. coli* UTI89 were selected for alignment. Proteins of interest from UTI89 are indicated with ***. The hypothetical protein (WP_000841005.1) is clustered with 17 others (A), and the hypothetical protein (WP_001298030.1) was clustered with 15 other similar proteins (C). (B and D) Protein sequences are aligned with Clustal Omega, and the matching amino acid residues are color-coded. The hypothetical protein from *E. coli* UTI89 (WP_000841005.1) is aligned with the proteins from *Salmonella enterica* (EBJ5110809.1) and *Yersinia entomophaga* (ABG33872) (B), and hypothetical protein (WP_001298030.1) is aligned with a protein form *S. enterica* (EAA7986694.1) and two other *E. coli* proteins (DksA, NP_414687.1 and YacG, NP_414643.1) (D). The residues in pink highlight the four cysteines coordinating the zinc finger feature (D). (E) Graphical representation of four *E. coli* species genomes with the homologues of UTI89_RS23725 and UTI89_RS27595 and their genomic location sites.

**Figure S4.**
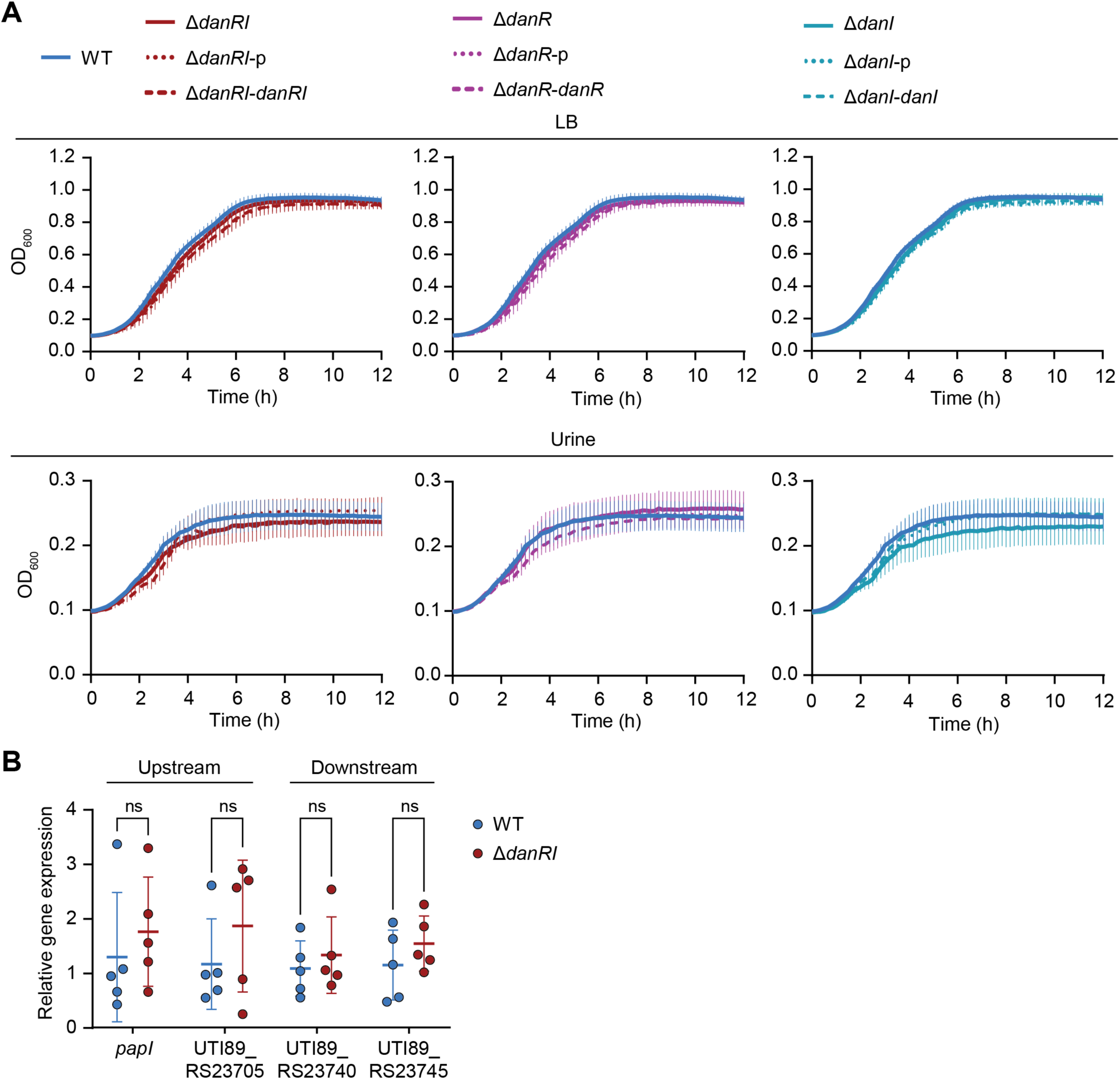
Mutants exhibit no growth defects or polar effects. (A) Wildtype UPEC, mutants, and their complemented strains were grown in LB medium or human urine for 12 hours while measuring the optical density at 600 nm (mean ± SEM, n = 3). (B) Gene expression measured by RT-qPCR of two upstream (*papI* and UTI89_RS23705) and two downstream (UTI89_RS23740 and UTI89-RS23745) genes in wildtype UPEC and Δ*danRI* relative to the housekeeping gene *gyrB* (mean ± SD, n = 5). Significance was determined with a Mann-Whitney test between wildtype and mutant strains (ns - no significant difference).

**Figure S5.**
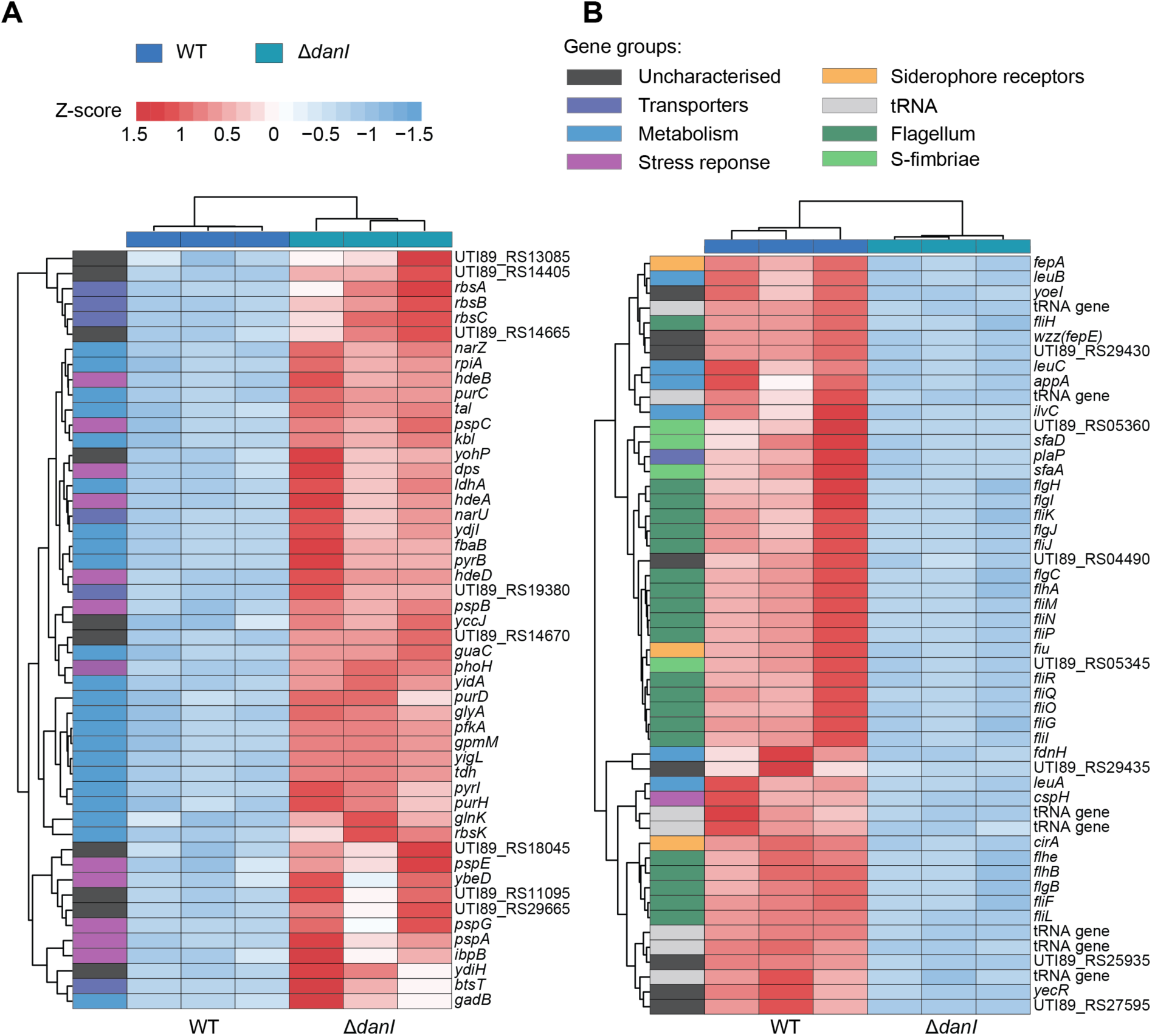
Δ*danI* exhibits a changed transcriptome. (A and B) Heatmap displaying Z scores of top 50 upregulated (A) and downregulated (B) genes in Δ*danI* compared to wildtype UPEC. The genes were clustered based on their normalized gene expression and were placed into categories based on their function: unknown/other, transporters, metabolism, stress response, siderophore receptors, tRNA, flagellum, and S-fimbriae.

**Figure S6.**
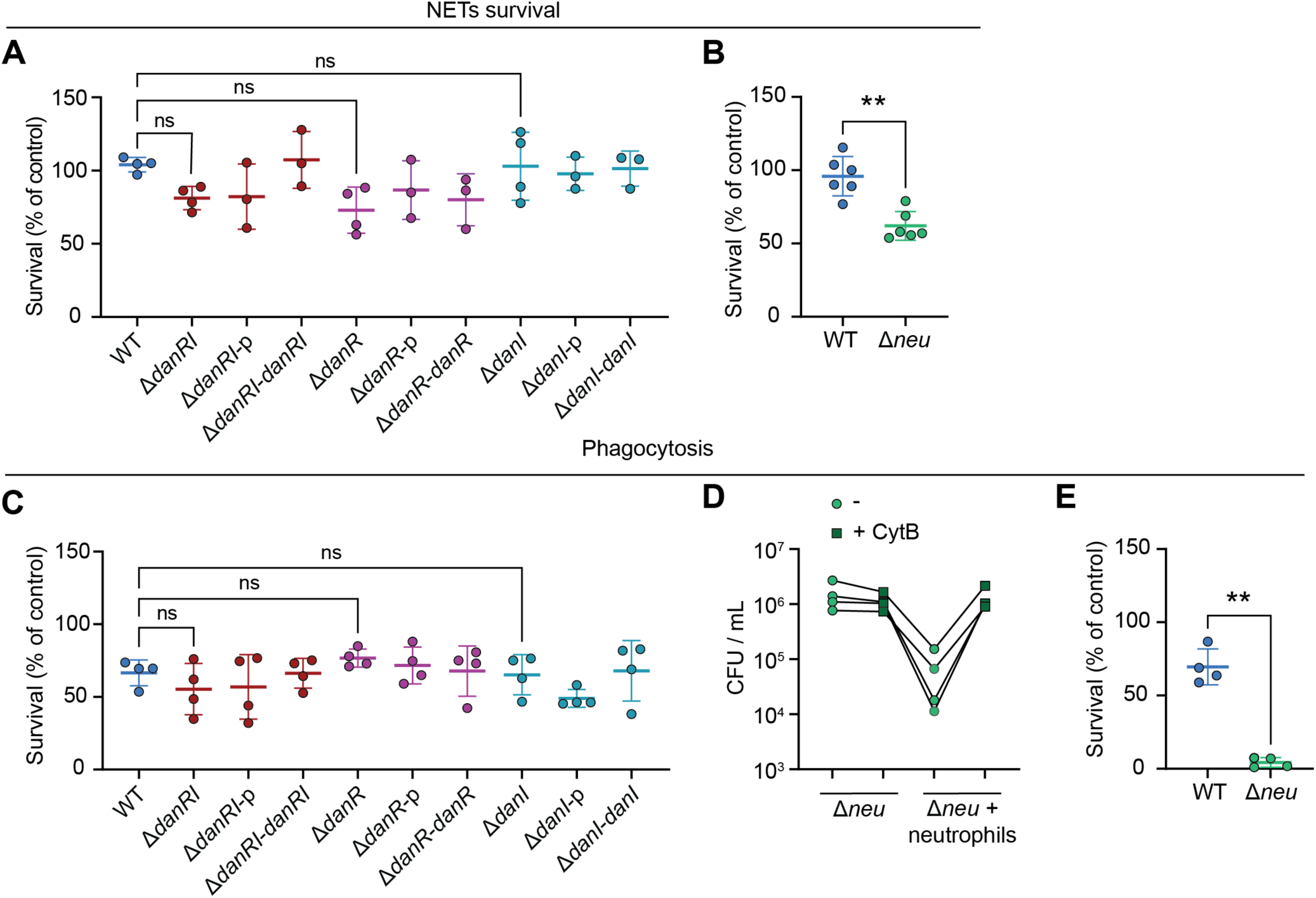
Capsule, but not DanRI, mutants are sensitive to phagosomal and NET killing. (A and B) Survival of wildtype UPEC, deletion mutants, and complemented strains on PMA-stimulated NETs for 1 hour is shown as a percentage of CFU on NETs normalized to CFU from medium with PMA (mean ± SD; A, n = 3-4; B, n = 6). (C, D, and E) Neutrophils with or without 10 µg/mL of Cytochalasin B were incubated with opsonized wildtype UPEC, deletion mutants, and their complemented strains for 30 minutes. The graphs show the bacterial survival as a percentage of CFU from lysed neutrophils normalized to CFU obtained from lysed cytochalasin B-inhibited neutrophils (mean ± SD; C - E, n = 4) or the resulting CFU/mL from such conditions connected by line matching the same experiment (D, n = 4). Significance was determined with a one-way ANOVA with Šídák’s multiple comparisons test (A and C, ns - no significant difference) and with a paired t-test (B and E, ** p ≤ 0.001).

**Figure S7.**
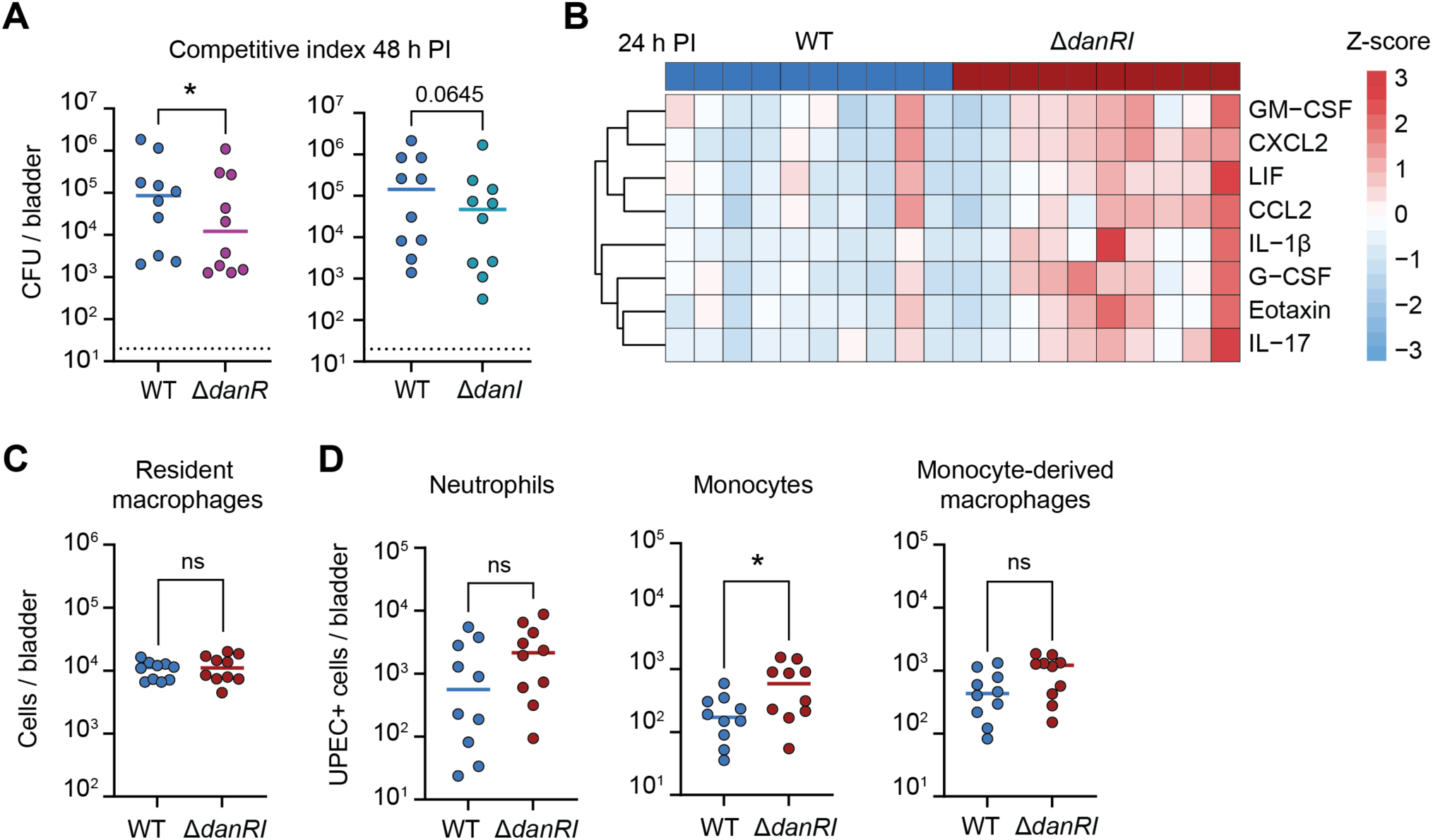
The regulatory system DanRI contributes to pathogenesis of UPEC. (A) CFU per bladder from infection with a 1:1 mixture of parental UPEC and Δ*danR* or Δ*danI* 48 hours post-infection (median, n = 10 mice from 2 independent experiments). Significance was determined with the Wilcoxon matched-pairs signed-rank test (* p ≤ 0.05). (B) Heatmap displaying Z-scores of GM-CSF, CXCL2, LIF, CCL2, IL-1β, G-CSF, eotaxin, and IL-17 concentrations from parental UPEC and Δ*danRI*-infected bladder at 24 hours post-infection (n = 10 mice for 2 independent experiments). (C) Total number of resident macrophages in parental UPEC and Δ*danRI*-infected bladders 24 hours post-infection (median from 10 mice from 2 independent experiments). (D) Number of neutrophils, monocytes, and monocyte-derived macrophages that are positive for UPEC expressing RFP in parental UPEC and Δ*danRI*-infected bladders 24 hours post-infection (median from 10 mice from 2 independent experiments). Significance was determined between parental UPEC and Δ*danRI* with the unpaired t-test corrected for the FDR (C and D, ns - no significant difference; * p ≤ 0.05).

### Supplemental tables

**Table S1.** Bacterial strains used in the study.

| Strain | Characteristics and function | Source and reference |
| --- | --- | --- |
| *E. coli* K12 | Wildtype commensal *E. coli* MG1655 K12 | Lab collection |
| UPEC | Wildtype *E. coli* UTI89<br>Uropathogenic strain isolated from a patient with cystitis | Lab collection<br>1 |
| EIEC | Wildtype enteroinvasive *E. coli* EIEC1 | Lab collection |
| EPEC | Wildtype enteropathogenic *E. coli* DEC5A | Lab collection |
| UPEC-RFP | *E. coli* UTI89 attB::Km-marsRFP; Km<sup>R</sup><br>Parental strain used in animal experiments, expressing RFP | 2 |
| UPEC-GFP | UTI89 attB::bla-GFP; Amp<sup>R</sup><br>Parental strain used in animal experiments, expressing GFP | 2 |
| ΔdanRI | *E. coli* UTI89ΔdanRI::FRT<br>Deletion mutant of the genomic region with *danR* and *danI* | This study |
| ΔdanRI-RFP | *E. coli* UTI89ΔdanRI::Km-marsRFP; Km<sup>R</sup><br>Deletion mutant of the genomic region with *danR* and *danI* expressing RFP, used for animal experiments | This study |
| ΔdanRI-p | *E. coli* UTI89ΔdanRI::FRT + pTrc99a; Amp<sup>R</sup><br>Deletion mutant of the genomic region with *danR* and *danI* with empty vector for complementation | This study |
| ΔdanRI-danRI | *E. coli* UTI89ΔdanRI::FRT + pTrc99a-danRI; Amp<sup>R</sup><br>Deletion mutant of the genomic region of *danR* and *danI* with a complementation vector carrying the genomic region | This study |
| ΔdanR | *E. coli* UTI89ΔdanR::FRT<br>Deletion mutant of *danR* | This study |
| ΔdanR-RFP | *E. coli* UTI89ΔdanR::Km-marsRFP; Km<sup>R</sup><br>Deletion mutant of *danR* expressing RFP, used for animal experiments | This study |
| ΔdanR-p | *E. coli* UTI89ΔdanR::FRT + pTrc99a; Amp<sup>R</sup><br>Deletion mutant of *danR* with empty vector for complementation | This study |
| ΔdanR-danR | *E. coli* UTI89ΔdanR::FRT + pTrc99a-danR; Amp<sup>R</sup><br>Deletion mutant of *danR* with a complementation vector carrying the *danR* | This study |
| ΔdanI | *E. coli* UTI89ΔdanI::FRT<br>Deletion mutant of *danI* | This study |
| ΔdanI-RFP | *E. coli* UTI89ΔdanI:: Km-marsRFP; Km<sup>R</sup><br>Deletion mutant of *danI* expressing RFP, used for animal experiments | This study |
| ΔdanI-p | *E. coli* UTI89ΔdanI::FRT + pTrc99a; Amp<sup>R</sup><br>Deletion mutant of *danI* with empty vector for complementation | This study |
| ΔdanI-danI | *E. coli* UTI89ΔdanI::FRT+ pTrc99a-danI; Amp<sup>R</sup><br>Deletion mutant of *danI* with a complementation vector carrying the *danI* | This study |
| UPEC-pGEN-*em7* | *E. coli* UTI89 + pGEN-*em7-luxCDABE*; Amp<sup>R</sup><br>With a vector expressing the luciferase operon under the *em7* constitutive promoter | This study |
| UPEC-pGEN | *E. coli* UTI89 + pGEN-*luxCDABE*; Amp<sup>R</sup><br>With a vector expressing the luciferase operon under no promoter | This study |
| UPEC-pGEN-prom-*danR* | *E. coli* UTI89 + pGEN-promoter-*danR-luxCDABE*; Amp<sup>R</sup><br>With a vector expressing the luciferase operon under the cloned *danR* promoter | This study |
| UPEC-pGEN-prom-*danI* | *E. coli* UTI89 + pGEN-promoter-*danI-luxCDABE*; Amp<sup>R</sup><br>With a vector expressing the luciferase operon under the cloned *danI* promoter | This study |

FRT = genome scar left by the Flippase Recognition Target site after recombination
Amp<sup>R</sup> = ampicillin/carbenicillin resistance
Km<sup>R</sup> = kanamycin resistance

**Table S2.**
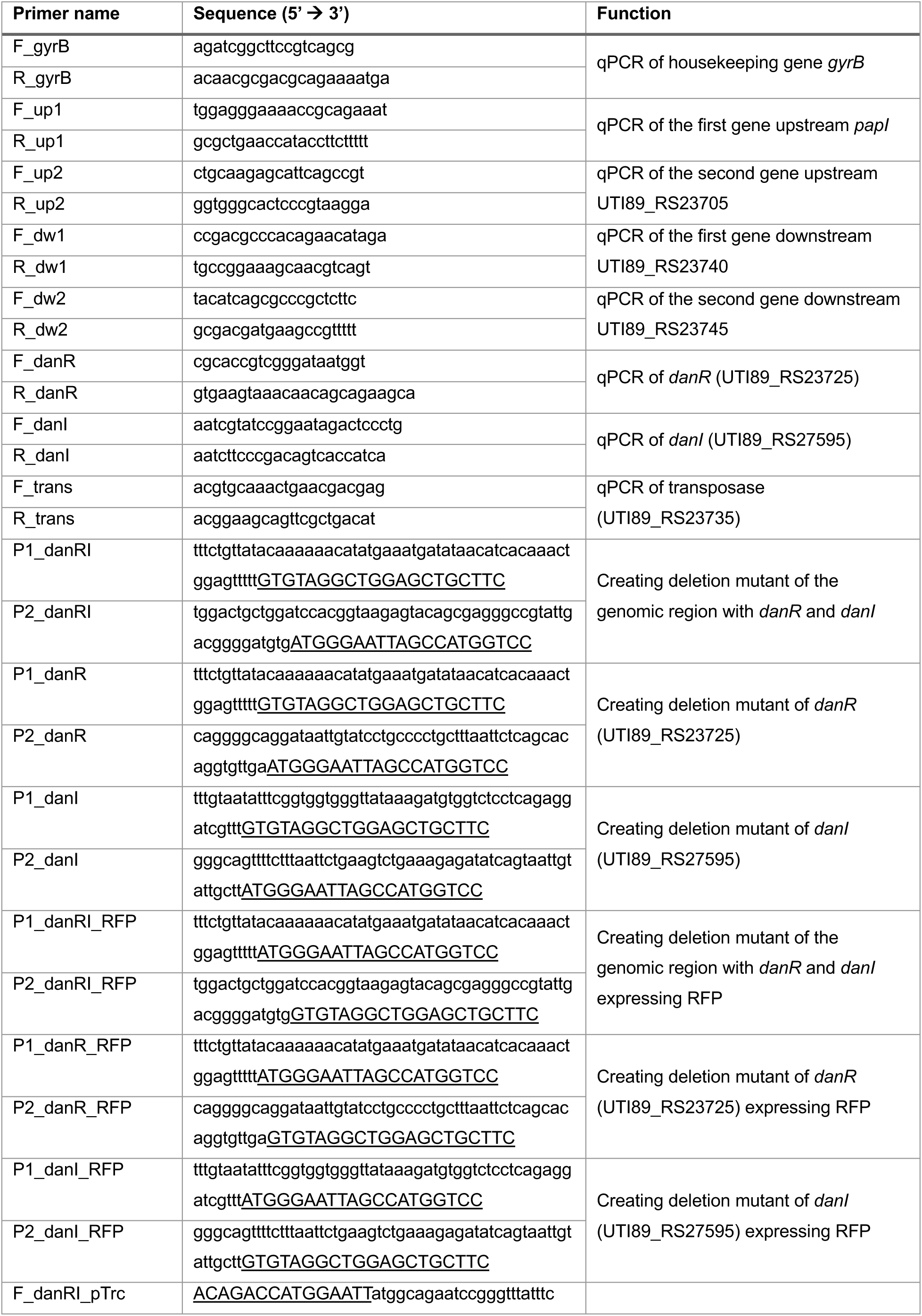
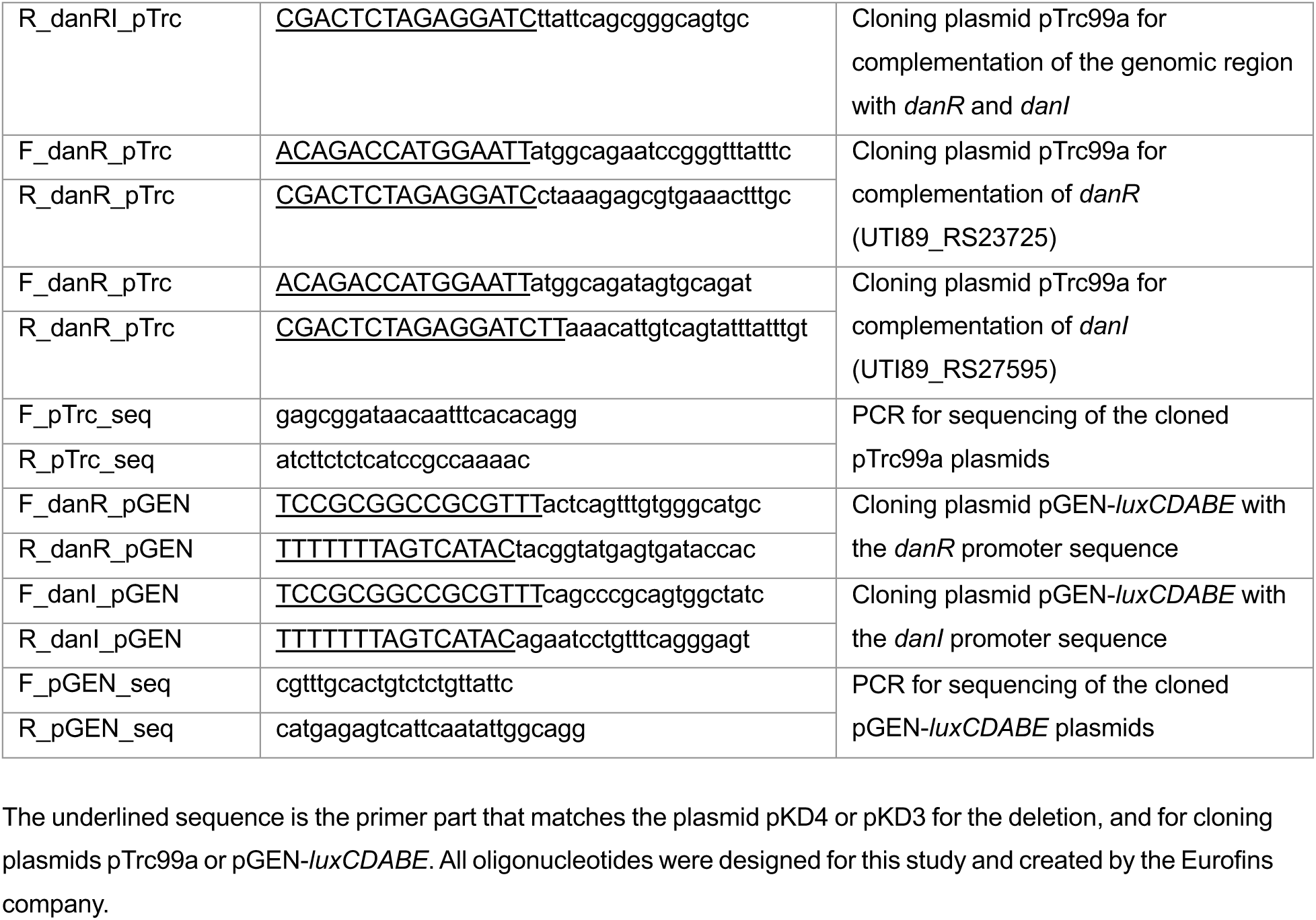
Oligonucleotides used in this study.

**Table S3.** Plasmids used in this study.

| Name | Characteristics | Source and reference |
| --- | --- | --- |
| pKM208 | Lambda Red recombination genes, *lacI* repression; Amp<sup>R</sup> | Lab collection |
| pKD4 | Template for kanamycin resistance genes flanked with FRT sites; Amp<sup>R</sup> | Lab collection |
| pKD3 | Template for chloramphenicol resistance genes flanked with FRT sites; Cm<sup>R</sup> | Lab collection |
| pCP20 | Thermal-induced FLP synthesis to remove FRT-flanked resistance gene; Amp<sup>R</sup> | Lab collection |
| pTrc99a | Trc promoter; Amp<sup>R</sup><br>Expression vector used for complementation | NovoPro Bioscience |
| pTrc99a-*danRI* | Expression vector used for complementation with the genomic region of *danR* and *danI*; Amp<sup>R</sup> | This study |
| pTrc99a-*danR* | Expression vector used for complementation with *danR*; Amp<sup>R</sup> | This study |
| pTrc99a-*danI* | Expression vector used for complementation with *danI*; Amp<sup>R</sup> | This study |
| pGEN-*em7-luxCDABE* | *Em7* promoter driving the luciferase operon; Amp<sup>R</sup> | Harry Mobley (Addgene # 44918)<sup>3</sup> |
| pGEN-*luxCDABE* | No promoter driving the luciferase operon; Amp<sup>R</sup> | This study |
| pGEN-promoter-*danR-luxCDABE* | *DanR* promoter sequence driving the luciferase operon; Amp<sup>R</sup> | This study |
| pGEN-promoter-*danI-luxCDABE* | *DanI* promoter sequence driving the luciferase operon; Amp<sup>R</sup> | This study |

**Table S4.** Antibodies used in this study to analyze bladder immune cells.

| **Antibody** | **Fluorochrome** | **Clone** | **Source and reference** |
| --- | --- | --- | --- |
| CX3CR1 | BV605 | SA011F11 | Biolegend |
| NK1.1 | Alexa Fluor 700 | PK136 | BD pharmingen |
| MHCII | APC-eFluor 780 | M5/114.15.2 | Invitrogen |
| F4/80 | Alexa Fluor 488 | Cl:A3-1 | Biorad |
| Ly6C | PerCP-Cy 5.5 | AL-21 | BD pharmingen |
| CD11b | BV421 | M1/70 | BD Horizon |
| CD103 | BV510 | M290 | BD Horizon |
| Ly6G | APC | 1A8 | BD pharmingen |
| CD64 | PE | X54-5/7.1 | BD pharmingen |
| SiglecF | BV786 | E50-2440 | BD optibuild |
| CD45 | BUV395 | 30-F11 | BD Horizon |
| CD62L | PE-Cy7 | MEL-14 | Invitrogen |
| CD11c | BV711 | HL3 | BD Horizon |

## References

1 Flores-Mireles, A. L., Walker, J. N., Caparon, M. & Hultgren, S. J. Urinary tract infections: epidemiology, mechanisms of infection and treatment options. Nat Rev Microbiol 13, 269–284, doi:10.1038/nrmicro3432 (2015).

2 Deltourbe, L., Lacerda Mariano, L., Hreha, T. N., Hunstad, D. A. & Ingersoll, M. A. The impact of biological sex on diseases of the urinary tract. Mucosal Immunol 15, 857–866, doi:10.1038/s41385-022-00549-0 (2022).

3 Ikähelmo, R. et al. Recurrence of Urinary Tract Infection in a Primary Care Setting: Analysis of a I-Year Follow-up of 179 Women. Clinical Infectious Diseases 22, 91–99, doi:10.1093/clinids/22.1.91 (1996).

4 Nicolas-Chanoine, M. H., Bertrand, X. & Madec, J. Y. *Escherichia coli* ST131, an intriguing clonal group. Clin Microbiol Rev 27, 543–574, doi:10.1128/cmr.00125-13 (2014).

5 Johnson, J. R., Johnston, B., Clabots, C., Kuskowski, M. A. & Castanheira, M. *Escherichia coli* sequence type ST131 as the major cause of serious multidrug-resistant *E. coli* infections in the United States. Clin Infect Dis 51, 286–294, doi:10.1086/653932 (2010).

6 Öztürk, R. & Murt, A. Epidemiology of urological infections: a global burden. World J Urol 38, 2669–2679, doi:10.1007/s00345-019-03071-4 (2020).

7 Kaper, J. B., Nataro, J. P. & Mobley, H. L. Pathogenic *Escherichia coli*. Nat Rev Microbiol 2, 123–140, doi:10.1038/nrmicro818 (2004).

8 Martinez, J. J., Mulvey, M. A., Schilling, J. D., Pinkner, J. S. & Hultgren, S. J. Type 1 pilus-mediated bacterial invasion of bladder epithelial cells. Embo j 19, 2803–2812, doi:10.1093/emboj/19.12.2803 (2000).

9 Lacerda Mariano, L. & Ingersoll, M. A. The immune response to infection in the bladder. Nat Rev Urol 17, 439–458, doi:10.1038/s41585-020-0350-8 (2020).

10 Ingersoll, M. A., Kline, K. A., Nielsen, H. V. & Hultgren, S. J. G-CSF induction early in uropathogenic *Escherichia coli* infection of the urinary tract modulates host immunity. Cell Microbiol 10, 2568–2578, doi:10.1111/j.1462-5822.2008.01230.x (2008).

11 Hannan, T. J. et al. Inhibition of Cyclooxygenase-2 Prevents Chronic and Recurrent Cystitis. EBioMedicine 1, 46–57, doi:10.1016/j.ebiom.2014.10.011 (2014).

12 Haraoka, M. et al. Neutrophil recruitment and resistance to urinary tract infection. J Infect Dis 180, 1220–1229, doi:10.1086/315006 (1999).

13 Cotzomi-Ortega, I. et al. Neutrophil NADPH oxidase promotes bacterial eradication and regulates NF-κB-Mediated inflammation via NRF2 signaling during urinary tract infections. Mucosal Immunology, doi:10.1016/j.mucimm.2024.12.010 (2024).

14 Burn, G. L., Foti, A., Marsman, G., Patel, D. F. & Zychlinsky, A. The Neutrophil. Immunity 54, 1377–1391, doi:10.1016/j.immuni.2021.06.006 (2021).

15 Brinkmann, V. et al. Neutrophil extracellular traps kill bacteria. Science 303, 1532–1535, doi:10.1126/science.1092385 (2004).

16 Yu, Y. et al. Diagnosing inflammation and infection in the urinary system via proteomics. J Transl Med 13, 111, doi:10.1186/s12967-015-0475-3 (2015).

17 Yu, Y. et al. Characterization of Early-Phase Neutrophil Extracellular Traps in Urinary Tract Infections. PLoS Pathog 13, e1006151, doi:10.1371/journal.ppat.1006151 (2017).

18 Krivošíková, K. et al. Neutrophil extracellular traps in urinary tract infection. Front Pediatr 11, 1154139, doi:10.3389/fped.2023.1154139 (2023).

19 Urban, C. F. et al. Neutrophil extracellular traps contain calprotectin, a cytosolic protein complex involved in host defense against *Candida albicans*. PLoS Pathog 5, e1000639, doi:10.1371/journal.ppat.1000639 (2009).

20 Marsman, G. et al. Histone H1 kills MRSA. Cell Rep 43, 114969, doi:10.1016/j.celrep.2024.114969 (2024).

21 Ganz, T. et al. Defensins. Natural peptide antibiotics of human neutrophils. J Clin Invest 76, 1427–1435, doi:10.1172/jci112120 (1985).

22 Parker, H., Albrett, A. M., Kettle, A. J. & Winterbourn, C. C. Myeloperoxidase associated with neutrophil extracellular traps is active and mediates bacterial killing in the presence of hydrogen peroxide. J Leukoc Biol 91, 369–376, doi:10.1189/jlb.0711387 (2012).

23 Fabisiak, A., Murawska, N. & Fichna, J. LL-37: Cathelicidin-related antimicrobial peptide with pleiotropic activity. Pharmacological Reports 68, 802–808, doi: 10.1016/j.pharep.2016.03.015 (2016).

24 Weinrauch, Y., Drujan, D., Shapiro, S. D., Weiss, J. & Zychlinsky, A. Neutrophil elastase targets virulence factors of enterobacteria. Nature 417, 91–94, doi:10.1038/417091a (2002).

25 Urban, C. F., Lourido, S. & Zychlinsky, A. How do microbes evade neutrophil killing? Cell Microbiol 8, 1687–1696, doi:10.1111/j.1462-5822.2006.00792.x (2006).

26 Pastorello, I. et al. EsiB, a novel pathogenic *Escherichia coli* secretory immunoglobulin A-binding protein impairing neutrophil activation. mBio 4, doi:10.1128/mBio.00206-13 (2013).

27 Lau, M. E., Loughman, J. A. & Hunstad, D. A. YbcL of uropathogenic *Escherichia coli* suppresses transepithelial neutrophil migration. Infect Immun 80, 4123–4132, doi:10.1128/iai.00801-12 (2012).

28 Ou, Q. et al. TcpC inhibits neutrophil extracellular trap formation by enhancing ubiquitination mediated degradation of peptidylarginine deiminase 4. Nat Commun 12, 3481, doi:10.1038/s41467-021-23881-8 (2021).

29 Breland, E. J., Eberly, A. R. & Hadjifrangiskou, M. An Overview of Two-Component Signal Transduction Systems Implicated in Extra-Intestinal Pathogenic *E. coli* Infections. Frontiers in Cellular and Infection Microbiology 7, doi:10.3389/fcimb.2017.00162 (2017).

30 Branzk, N. et al. Neutrophils sense microbe size and selectively release neutrophil extracellular traps in response to large pathogens. Nat Immunol 15, 1017–1025, doi:10.1038/ni.2987 (2014).

31 Amulic, B. et al. Cell-Cycle Proteins Control Production of Neutrophil Extracellular Traps. Dev Cell 43, 449–462.e445, doi:10.1016/j.devcel.2017.10.013 (2017).

32 Kenny, E. F. et al. Diverse stimuli engage different neutrophil extracellular trap pathways. Elife 6, doi:10.7554/eLife.24437 (2017).

33 Campbell, E. A. et al. Structural mechanism for rifampicin inhibition of bacterial rna polymerase. Cell 104, 901–912, doi:10.1016/s0092-8674(01)00286-0 (2001).

34 Ashburner, M. et al. Gene ontology: tool for the unification of biology. The Gene Ontology Consortium. Nat Genet 25, 25–29, doi:10.1038/75556 (2000).

35 Sidote, D. J., Barbieri, C. M., Wu, T. & Stock, A. M. Structure of the *Staphylococcus aureus* AgrA LytTR domain bound to DNA reveals a beta fold with an unusual mode of binding. Structure 16, 727–735, doi:10.1016/j.str.2008.02.011 (2008).

36 McGowan, S. et al. The FxRxHrS motif: a conserved region essential for DNA binding of the VirR response regulator from *Clostridium perfringens*. J Mol Biol 322, 997–1011, doi:10.1016/s0022-2836(02)00850-1 (2002).

37 Chen, S. L. et al. Identification of genes subject to positive selection in uropathogenic strains of *Escherichia coli:* a comparative genomics approach. Proc Natl Acad Sci U S A 103, 5977–5982, doi:10.1073/pnas.0600938103 (2006).

38 Desvaux, M. et al. Pathogenicity Factors of Genomic Islands in Intestinal and Extraintestinal *Escherichia coli*. Front Microbiol 11, 2065, doi:10.3389/fmicb.2020.02065 (2020).

39 Johnson, M. et al. NCBI BLAST: a better web interface. Nucleic Acids Res 36, W5–9, doi:10.1093/nar/gkn201 (2008).

40 van Kempen, M. et al. Fast and accurate protein structure search with Foldseek. Nature Biotechnology 42, 243–246, doi:10.1038/s41587-023-01773-0 (2024).

41 Perederina, A. et al. Regulation through the secondary channel--structural framework for ppGpp-DksA synergism during transcription. Cell 118, 297–309, doi:10.1016/j.cell.2004.06.030 (2004).

42 Sengupta, S. & Nagaraja, V. YacG from *Escherichia coli* is a specific endogenous inhibitor of DNA gyrase. Nucleic Acids Research 36, 4310–4316, doi:10.1093/nar/gkn355 (2008).

43 Ramelot, T. A. et al. NMR structure of the *Escherichia coli* protein YacG: A novel sequence motif in the zinc-finger family of proteins. *Proteins: Structure*, Function, and Bioinformatics 49, 289–293, doi:10.1002/prot.10214 (2002).

44 Henard, C. A. et al. The 4-cysteine zinc-finger motif of the RNA polymerase regulator DksA serves as a thiol switch for sensing oxidative and nitrosative stress. Mol Microbiol 91, 790–804, doi:10.1111/mmi.12498 (2014).

45 Jolley, K. A., Bray, J. E. & Maiden, M. C. J. Open-access bacterial population genomics: BIGSdb software, the PubMLST.org website and their applications. Wellcome Open Res 3, 124, doi:10.12688/wellcomeopenres.14826.1 (2018).

46 Denamur, E., Clermont, O., Bonacorsi, S. & Gordon, D. The population genetics of pathogenic *Escherichia coli*. Nat Rev Microbiol 19, 37–54, doi:10.1038/s41579-020-0416-x (2021).

47 Emery, A., Hocquet, D., Bonnet, R. & Bertrand, X. Genotypic Characteristics and Antimicrobial Resistance of *Escherichia coli* ST141 Clonal Group. Antibiotics (Basel) 12, 382, doi:10.3390/antibiotics12020382 (2023).

48 Cave, R., Ter-Stepanyan, M. M., Kotsinyan, N. & Mkrtchyan, H. V. An Emerging Lineage of Uropathogenic Extended Spectrum β-Lactamase *Escherichia coli* ST127. Microbiology Spectrum 10, e02511–02522, doi:10.1128/spectrum.02511-22 (2022).

49 Horner, C., et al. *Escherichia coli* bacteraemia: 2 years of prospective regional surveillance (2010–12). Journal of Antimicrobial Chemotherapy 69, 91–100, doi:10.1093/jac/dkt333 (2014).

50 Salgado, H., Moreno-Hagelsieb, G., Smith, T. F. & Collado-Vides, J. Operons in *Escherichia coli:* genomic analyses and predictions. Proc Natl Acad Sci U S A 97, 6652–6657, doi:10.1073/pnas.110147297 (2000).

51 Zhang, H. et al. Metabolic stress promotes stop-codon readthrough and phenotypic heterogeneity. Proc Natl Acad Sci U S A 117, 22167–22172, doi:10.1073/pnas.2013543117 (2020).

52 Datsenko, K. A. & Wanner, B. L. One-step inactivation of chromosomal genes in *Escherichia coli* K-12 using PCR products. Proc Natl Acad Sci U S A 97, 6640–6645, doi:10.1073/pnas.120163297 (2000).

53 Murphy, K. C. & Campellone, K. G. Lambda Red-mediated recombinogenic engineering of enterohemorrhagic and enteropathogenic *E. coli*. BMC Mol Biol 4, 11, doi:10.1186/1471-2199-4-11 (2003).

54 Anderson, G. G., Goller, C. C., Justice, S., Hultgren, S. J. & Seed, P. C. Polysaccharide capsule and sialic acid-mediated regulation promote biofilm-like intracellular bacterial communities during cystitis. Infect Immun 78, 963–975, doi:10.1128/iai.00925-09 (2010).

55 Wartha, F. et al. Capsule and D-alanylated lipoteichoic acids protect *Streptococcus pneumoniae* against neutrophil extracellular traps. Cell Microbiol 9, 1162–1171, doi:10.1111/j.1462-5822.2006.00857.x (2007).

56 Li, Y. & Trush, M. A. Diphenyleneiodonium, an NAD(P)H oxidase inhibitor, also potently inhibits mitochondrial reactive oxygen species production. Biochem Biophys Res Commun 253, 295–299, doi:10.1006/bbrc.1998.9729 (1998).

57 Tilley, D. O. et al. Histone H3 clipping is a novel signature of human neutrophil extracellular traps. Elife 11, doi:10.7554/eLife.68283 (2022).

58 Zychlinsky Scharff, A., Albert, M. L. & Ingersoll, M. A. Urinary Tract Infection in a Small Animal Model: Transurethral Catheterization of Male and Female Mice. J Vis Exp, doi:10.3791/54432 (2017).

59 Mora-Bau, G. et al. Macrophages Subvert Adaptive Immunity to Urinary Tract Infection. PLoS Pathog 11, e1005044, doi:10.1371/journal.ppat.1005044 (2015).

60 Zychlinsky Scharff, A., et al. Sex differences in IL-17 contribute to chronicity in male versus female urinary tract infection. JCI Insight 5, doi:10.1172/jci.insight.122998 (2019).

61 Loughman, J. A. & Hunstad, D. A. Attenuation of human neutrophil migration and function by uropathogenic bacteria. Microbes Infect 13, 555–565, doi:10.1016/j.micinf.2011.01.017 (2011).

62 Lv, H., Hung, C. S. & Henderson, J. P. Metabolomic analysis of siderophore cheater mutants reveals metabolic costs of expression in uropathogenic *Escherichia coli*. J Proteome Res 13, 1397–1404, doi:10.1021/pr4009749 (2014).

63 Okinaga, T., Niu, G., Xie, Z., Qi, F. & Merritt, J. The hdrRM operon of *Streptococcus mutans* encodes a novel regulatory system for coordinated competence development and bacteriocin production. J Bacteriol 192, 1844–1852, doi:10.1128/jb.01667-09 (2010).

64 Xie, Z., Okinaga, T., Niu, G., Qi, F. & Merritt, J. Identification of a novel bacteriocin regulatory system in *Streptococcus mutans*. Molecular Microbiology 78, 1431–1447, doi:10.1111/j.1365-2958.2010.07417.x (2010).

65 Zou, Z. et al. LytTR Regulatory Systems: A potential new class of prokaryotic sensory system. PLOS Genetics 14, e1007709, doi:10.1371/journal.pgen.1007709 (2018).

66 Cheung, J. K. & Rood, J. I. The VirR response regulator from *Clostridium perfringens* binds independently to two imperfect direct repeats located upstream of the pfoA promoter. J Bacteriol 182, 57–66, doi:10.1128/jb.182.1.57-66.2000 (2000).

67 Nikolskaya, A. N. & Galperin, M. Y. A novel type of conserved DNA-binding domain in the transcriptional regulators of the AlgR/AgrA/LytR family. Nucleic Acids Res 30, 2453–2459, doi:10.1093/nar/30.11.2453 (2002).

68 Matthews, J. M. & Sunde, M. Zinc Fingers-Folds for Many Occasions. IUBMB Life 54, 351–355, doi:10.1080/15216540216035 (2002).

69 Cai, W. et al. A novel two-component signaling system facilitates uropathogenic *Escherichia coli*’s ability to exploit abundant host metabolites. PLoS Pathog 9, e1003428, doi:10.1371/journal.ppat.1003428 (2013).

70 Gu, H. et al. A previously uncharacterized two-component signaling system in uropathogenic *Escherichia coli* coordinates protection against host-derived oxidative stress with activation of hemolysin-mediated host cell pyroptosis. PLoS Pathog 17, e1010005, doi:10.1371/journal.ppat.1010005 (2021).

71 Li, X. et al. Two-component system GrpP/GrpQ promotes pathogenicity of uropathogenic *Escherichia coli* CFT073 by upregulating type 1 fimbria. Nature Communications 16, 607, doi:10.1038/s41467-025-55982-z (2025).

72 Sultana, S. et al. Redox-Mediated Inactivation of the Transcriptional Repressor RcrR is Responsible for Uropathogenic *Escherichia coli*’s Increased Resistance to Reactive Chlorine Species. mBio 13, e0192622, doi:10.1128/mbio.01926-22 (2022).

73 Nguyen, G. T., Green, E. R. & Mecsas, J. Neutrophils to the ROScue: Mechanisms of NADPH Oxidase Activation and Bacterial Resistance. Frontiers in Cellular and Infection Microbiology 7, doi:10.3389/fcimb.2017.00373 (2017).

74 De Filippo, K. et al. Mast cell and macrophage chemokines CXCL1/CXCL2 control the early stage of neutrophil recruitment during tissue inflammation. Blood 121, 4930–4937, doi:10.1182/blood-2013-02-486217 (2013).

75 Hannan, T. J., Mysorekar, I. U., Hung, C. S., Isaacson-Schmid, M. L. & Hultgren, S. J. Early severe inflammatory responses to uropathogenic *E. coli* predispose to chronic and recurrent urinary tract infection. PLoS Pathog 6, e1001042, doi:10.1371/journal.ppat.1001042 (2010).

76 Veldhoen, M. Interleukin 17 is a chief orchestrator of immunity. Nature Immunology 18, 612–621, doi:10.1038/ni.3742 (2017).

77 Flannigan, K. L. et al. IL-17A-mediated neutrophil recruitment limits expansion of segmented filamentous bacteria. Mucosal Immunology 10, 673–684, doi:10.1038/mi.2016.80 (2017).

78 Patel, D. F. et al. Neutrophils restrain allergic airway inflammation by limiting ILC2 function and monocyte-dendritic cell antigen presentation. Sci Immunol 4, doi:10.1126/sciimmunol.aax7006 (2019).

79 Yamanaka, Y., Aizawa, S.-I. & Yamamoto, K. The *hdeD* Gene Represses the Expression of Flagellum Biosynthesis via LrhA in *Escherichia coli* K-12. Journal of Bacteriology 204, e00420–00421, doi:10.1128/JB.00420-21 (2022).

80 Tschowri, N., Busse, S. & Hengge, R. The BLUF-EAL protein YcgF acts as a direct anti-repressor in a blue-light response of *Escherichia coli*. Genes Dev 23, 522–534, doi:10.1101/gad.499409 (2009).

81 Jhelum, H. et al. Panton-Valentine leukocidin-induced neutrophil extracellular traps lack antimicrobial activity and are readily induced in patients with recurrent PVL+-*Staphylococcus aureus* infections. J Leukoc Biol, doi:10.1093/jleuko/qiad137 (2023).

82 Langmead, B. & Salzberg, S. L. Fast gapped-read alignment with Bowtie 2. Nature Methods 9, 357–359, doi:10.1038/nmeth.1923 (2012).

83 Liao, Y., Smyth, G. K. & Shi, W. featureCounts: an efficient general purpose program for assigning sequence reads to genomic features. Bioinformatics 30, 923–930, doi:10.1093/bioinformatics/btt656 (2013).

84 Love, M. I., Huber, W. & Anders, S. Moderated estimation of fold change and dispersion for RNA-seq data with DESeq2. Genome Biology 15, 550, doi:10.1186/s13059-014-0550-8 (2014).

85 Zhu, A., Ibrahim, J. G. & Love, M. I. Heavy-tailed prior distributions for sequence count data: removing the noise and preserving large differences. Bioinformatics 35, 2084–2092, doi:10.1093/bioinformatics/bty895 (2018).

86 Madeira, F. et al. The EMBL-EBI Job Dispatcher sequence analysis tools framework in 2024. Nucleic acids research 52, W521–W525, doi:10.1093/nar/gkae241 (2024).

87 Yamanaka, Y., Watanabe, H., Yamauchi, E., Miyake, Y. & Yamamoto, K. Measurement of the Promoter Activity in *Escherichia coli* by Using a Luciferase Reporter. Bio Protoc 10, e3500, doi:10.21769/BioProtoc.3500 (2020).

88 Schindelin, J. et al. Fiji: an open-source platform for biological-image analysis. Nature Methods 9, 676–682, doi:10.1038/nmeth.2019 (2012).

## References

1 Mulvey, M. A., Schilling, J. D. & Hultgren, S. J. Establishment of a persistent *Escherichia coli r*eservoir during the acute phase of a bladder infection. Infect Immun 69, 4572–4579, doi:10.1128/iai.69.7.4572-4579.2001 (2001).

2 Mora-Bau, G. et al. Macrophages Subvert Adaptive Immunity to Urinary Tract Infection. PLoS Pathog 11, e1005044, doi:10.1371/journal.ppat.1005044 (2015).

3 Lane, M. C., Alteri, C. J., Smith, S. N. & Mobley, H. L. Expression of flagella is coincident with uropathogenic *Escherichia coli* ascension to the upper urinary tract. Proc Natl Acad Sci U S A 104, 16669–16674, doi:10.1073/pnas.0607898104 (2007).

